# Determination of α-Synuclein Protein Interactions by μMap Photo-proximity Labeling

**DOI:** 10.1101/2025.09.18.674129

**Authors:** Marshall G. Lougee, Grace S. H. Park, Hee Jong Kim, Jiya Z. Fowler, Zongtao Lin, Hossein Fazelinia, Lynn Spruce, Evan Yanagawa, Benjamin A. Garcia, Melike Lakadamyali, E. James Petersson

## Abstract

Fibrillar aggregates of the natively disordered protein α-synuclein (αS) are hallmarks of Parkinson’s disease and related neurodegenerative disorders termed synucleinopathies. Here, we used micromap (µMap) photo-proximity labeling to determine the interactomes of αS monomers and fibrils in mouse brain lysate to better understand both the loss of healthy function and gain of toxic function aspects of synucleinopathies. Several αS variants were synthesized and characterized, showing that the small size (1 kDa) of the Ir catalyst attached through a Cys-maleimide linkage makes it minimally-perturbing to αS, with a narrow labeling radius that allows one to identify interactome differences between different regions of αS. Monomer and fibril interactomes were compared to each other and to previous proximity labeling data sets for validation and several examples of further investigations are demonstrated, including Western blotting, super-resolution microscopy, and µMap in primary neurons.

## Introduction

Identifying protein interactions is pivotal for a deeper understanding of biological function. A powerful way of ascertaining this information is proximity labeling (PL), which involves fusing a protein-of-interest (POI) with a labeling system capable of covalently modifying nearby interactors, particularly with affinity purification handles like biotin. These interacting molecules, which are often other proteins, can then be enriched by streptavidin/biotin pulldown and analyzed using mass spectrometry (MS) proteomic workflows.^1-8^ Numerous technologies exist for achieving this goal that take advantage of a genetically incorporated POI-enzyme fusion utilizing biotin ligase (BioID^9-10^/TurboID/miniTurbo^11-13^), ascorbate peroxidase (APEX1^14-15^/APEX2^16-17^), or a modified light-dependent enzyme (LITag^18^/miniSOG^19^). With each generation of these technologies, an emerging trend has been optimizing the labeling parameters to reduce perturbation to the native system while also highlighting the unique benefits of each PL platform.^20-22^ In this sense, there is not a single PL technology that should be utilized over all others, but rather the benefits and drawbacks should be weighed based on the scientific hypothesis to be tested.

An alternative to enzymatic fusion was presented in 2020 by the MacMillan lab.^23^ This novel PL platform instead utilizes an Ir photocatalyst attached to a POI paired with an accompanying biotin-diazirine (Dz-BTN) probe molecule (**Figure 1A-1B**). Upon irradiation with blue light, the Ir catalyst can transfer energy to the nearby Dz-BTN, which then extrudes N_2_, generating a reactive carbene capable of covalently labeling nearby proteins or biomolecules. Due to the strict distance constraints of the energy transfer process (≤10 Å) as well as the very short lifetime of the singlet carbene (∼2 ns), biotinylation is tightly clustered around the interacting area. This technology was termed “micromap” (µMap) due to the unprecedented small labeling (∼3-6 nm theoretical) radius versus other PL platforms and has since shown utility in multiple instances including immune cell profiling^23^, small molecule interactome profiling,^24^ SARS-CoV-2 entry mechanisms,^25^ e9 glycoprotein interactions,^26^ and chromatin interactome dynamics.^27^ Furthermore, red-shifted variants have been developed using either Sn or Os photocatalysts^28-29^ as well as a tunable variant using the organic photocatalyst Eosin Y termed “multimap.”^30^ Although photocatalytic systems such as µMap have smaller probe sizes (1 kDa) and labeling radii than enzymatic variants, there has thus far been limited exploration of direct protein attachment to truly exploit these features.^31^

**Figure 1.**
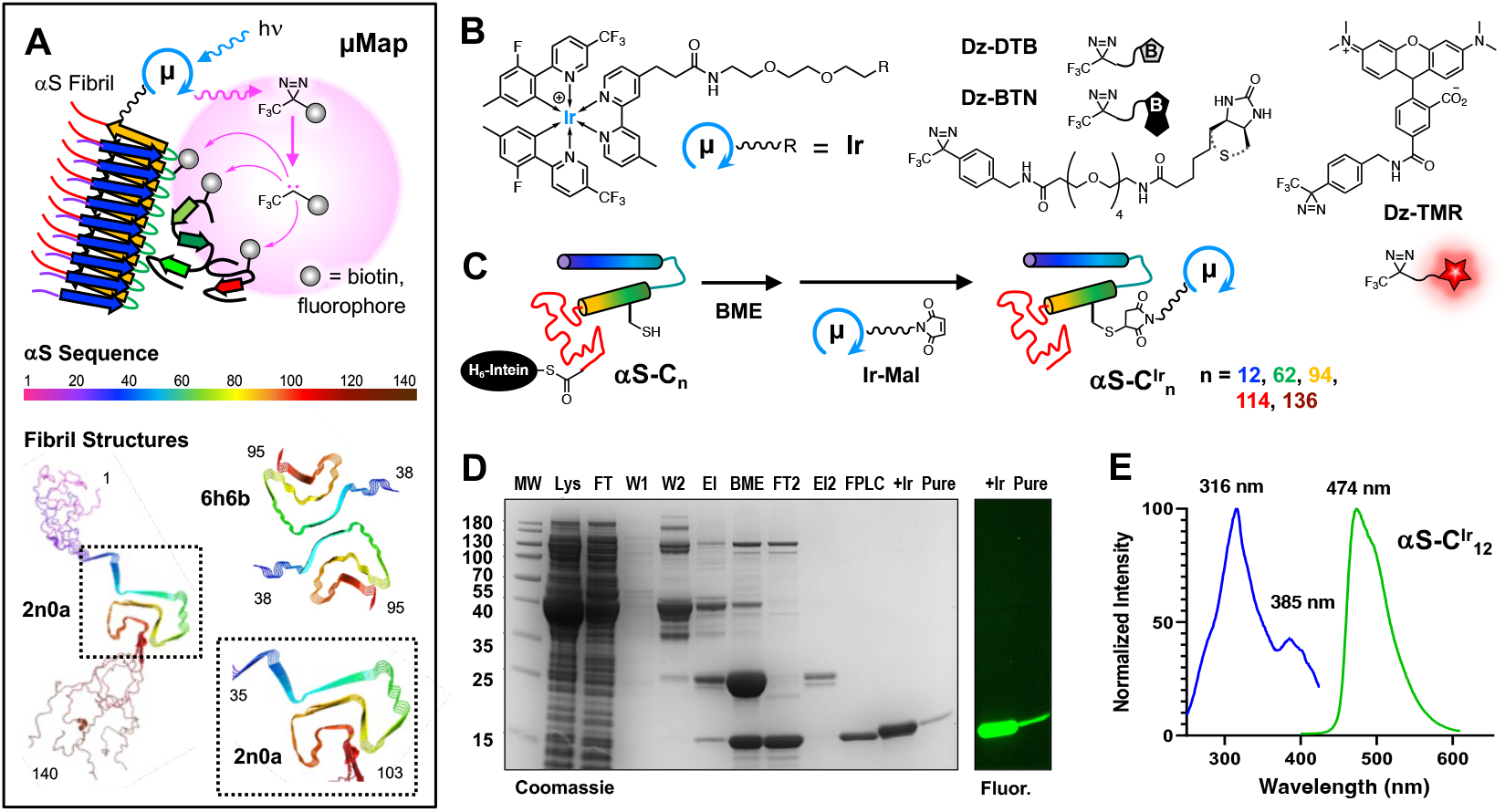
αS µMap Strategy and Protein Modification. (A) Top: In µMap proximity labelling a photocatalyst absorbs visible light and transfers energy to a freely-difusing diazirine probe, generating a high local concentration of carbenes that react with nearby proteins, labelling them with fluorophores for imaging or biotin for purification and identification through MS. Bottom: αS sequence and two representative fibril structures from ssNMR (PDB ID 2n0a)^32^ and cryo-EM (PDB ID 6h6b),^33^ colored using the same scale. (B) Structures and icons for iridium catalyst (Ir), desthiobiotin (Dz-DTB, without dashed thioether), biotin (Dz-BTN, with dashed thioether), or tetramethylrhodamine (Dz-TMR) diazirines. (C) Expression and Ir-maleimide (Ir-Mal) labelling scheme for αS-C^Ir^_n_ constructs. (D) Coomassie-stained gel and fluorescence gel section showing αS-C^Ir^_12_ purification and Ir-Mal labelling. (E) Absorption and emission spectra for αS-C^Ir^_12_ in PBS buffer.

Here, we investigated a variant of the µMap system to determine protein-protein interactions for a protein system that could adopt multiple conformations, the small intrinsically disordered protein α-synuclein (αS). The native function of αS is still not entirely clear and it has been reported to participate in a wide variety of biological processes, with a correspondingly wide array of protein interactions. αS is perhaps best known for its role in Parkinson’s disease (PD), where it forms amyloid-type fibrils that are part of pathological inclusions called Lewy bodies or Lewy neurites.^34-36^ These inclusions are thought to be both a result of upstream neurodegeneration and to contribute directly to further neuronal stress. Therefore, the loss of healthy αS interactions as well as the gain of pathological αS interactions may contribute to PD. Thus, the protein interactomes of both the monomeric and fibrillar forms of this protein are of great interest.

As a consequence of the interest in its healthy and PD interactomes, αS has been extensively studied in PL experiments. Chung *et* al. utilized the peroxidase-based APEX2 system to examine the interactome of monomeric αS in both HEK-293 cells and rat cortical neurons, identifying several key interactors including MAPT, DNAJ6C, and SYNJ1.^37^ In 2022, Killinger *et al*. followed up this initial study with *in situ* labeling of Lewy body interactors using an αS antibody paired with a secondary antibody fused to a peroxidase.^38^ BioID has also been used to assess the αS interactome by Griffin *et al*., who used differentiated embryonic stem cells paired with an αS-BioID2 fusion to assess the interactome of both monomeric and fibrillar αS.^39^ However, µMap offers a unique opportunity to interrogate the positional dependence of the αS interactome and compare the results to studies using these other PL technologies.

The Ir catalyst is expected to be less perturbing to protein folding and protein-protein interactions than a ∼20-30 kDa protein fusion (BioID/APEX). This is particularly important since it is known that point mutations and post-translational modifications can have a dramatic impact on αS folding.^40^ The theoretical labeling radius of the carbene is also smaller than the activated ester of BioID or the phenoxy radical of APEX, yielding a tighter interactome map.^5^ Another appealing characteristic is that the catalyst can be site-specifically attached to αS at a sidechain, not limited to N-terminal or C-terminal fusions of labeling proteins. The combination of variable attachment points and the tight labeling radius could lead to the generation of site-specific interactomes; identifying not only which proteins are interacting, but also with what portion of αS they are interacting **(Figure 1A)**.

## Results and Discussion

### µMap Probe Development and αS Labeling Optimization

To identify sites for attachment of the Ir catalyst that would not be expected to disrupt fibril formation, we consulted the available cryo-electron microscopy (cryo-EM) and solid state NMR (ssNMR) structures of αS fibrils made under conditions similar to those used in our laboratory.^32-33^ We chose locations in the N-terminus (1-60), the NAC-domain (61-95), and the C-terminus (96-140): lysine 12, glutamine 62, phenylalanine 94, glutamate 114, and tyrosine 136. These are sites that are either in the disordered regions of the fibrils or on the exterior faces of the folded core region (**Figure 1A**). They are also positions that we have previously determined to tolerate labeling with fluorescent probes similar in size to the Ir catalyst.^41-43^

In previous µMap studies, the MacMillan group utilized a dibenzocyclooctyne (DBCO) strain-promoted click reaction to attach their iridium catalyst to an antibody. With this in mind, we postulated that by using a DBCO-functionalized catalyst paired with azido-phenylalanine (AzF) unnatural amino acid mutagenesis, we would be able to incorporate the µMap system site-specifically in αS. As an alternative to this, we could also take advantage of the lack of cysteine residues in αS. By site specifically mutating cysteine in at various positions, we rationalized that a maleimide functionalized catalyst would also enable site-specific αS functionalization. A small array of accompanying diazirine probe molecules was synthesized, some of which have not been used in µMap labeling platforms before. These include a tetramethyl rhodamine diazirine derivative (Dz-TMR) for fluorescent labeling, a desthiobiotin derivative (Dz-DTB) for elution from streptavidin and subsequent crosslinking site identification, and the standard biotin derivative (Dz-BTN) for enhanced chemiluminescent (ECL) studies and standard proteomic pulldown experiments (**Figure 1B**).

αS AzF mutants were obtained by introducing the amber stop codon (TAG) in an αS-intein fusion vector for use in a traceless intein-based purification strategy described previously in our lab.^44^ Amber suppression was then conducted and mutants with AzF at phenylalanine 94 (αS-Z_94_), glutamate 114 (αS-Z_114_), and glutamate 136 (αS-Z_136_) were isolated. Initial labeling studies were conducted using a TAMRA-DBCO probe to assess the efficiency of the click reaction in AzF mutants. Unfortunately, poor labeling efficiency was observed for all mutants with no construct proceeding past ∼50% conversion by matrix-assisted laser desorption ionization mass spectrometry (MALDI-MS) analysis (See Supporting Information, SI, **Figure S1**). This factor compounded by low expression yields (0.51 mg protein per L culture) from unnatural amino acid mutagenesis, led us to pursue a different strategy in attaching the µMap catalyst to αS. αS contains no native cysteines, so insertion of an exogenous Cys residue followed by reaction with a maleimide affords a site-specifically labeled protein. Five αS cysteine mutants were expressed and purified using a similar intein-based purification strategy (**Figure 1C**), to generate αS-C_n_, n = 12, 62, 94, 114, or 136 in yields of 2.71-5.75 mg/L of initial culture.

To assess labeling efficiency, a new variant of iridium catalyst was synthesized that contained maleimide (Ir-Mal) for conjugation rather than DBCO (**Figure 1C**). Initial labeling studies showed improvement versus previously attempted DBCO labeling, and rigorous optimization led to a set of conditions resulting in quantitative conversion to iridium labeled αS (SI, **Figure S2**). Interestingly, decreasing both the tris(2-carboxyethyl) phosphine (TCEP) reducing agent and αS concentrations during labeling were crucial for optimal conversion to labeled protein. Excess Ir-Mal photocatalyst was removed via high performance liquid chromatography (HPLC) purification, and the five Ir-labeled αS positional variants, αS-C^Ir^_n_, n = 12, 62, 94, 114, or 136, were obtained in yields of 0.74-2.02 mg/L after labeling. Successful purification and labeling were determined by MALDIMS (SI, **Figure S3**), Coomassie gel analysis, and fluorescence analysis (**Figure 1D**), as the iridium catalyst itself shows green wavelength fluorescence (λ_max, exc._ = 316 nm; λ_max, em_. = 474 nm) (**Figure 1E**). We anticipate that this characteristic may be exploited in the future for potential imaging applications.

To assess the activity of our αS µMap system, Dz-TMR was synthesized via conjugation of trifluoromethyl diazirine phenyl methanamine with a TAMRA succinimidyl ester in DMF (See SI for synthesis). A preliminary set of *in vitro* benchmarking experiments were conducted with monomeric αS-C^Ir^_12_ in the presence of Dz-TMR as a measure of fluorescent self-labeling (SI **Figure S4**). Samples containing αS-C^Ir^_12_ and Dz-TMR were irradiated using a 456 nm Kessil PR160L lamp for 30 minutes with shaking. Rewardingly, all controls without Dz-TMR probe or light showed no fluorescence by gel analysis. Boiling of samples prior to gel analysis did not lead to any unanticipated side reactions, and diazirines did not show any more labeling in ambient light versus complete darkness; indicating that aluminum foil and/or excessively careful handling is unnecessary in these types of studies. Upon irradiation, samples containing Dz-TMR showed TAMRA fluorescence signal on the gel, indicating the expected self-labeling of the αS-C^Ir^_12_ monomer. Iridium fluorescence (green signal) was also observed for all samples and was overlayed with the TAMRA signal. Interestingly, a slight decrease in iridium fluorescence is observed in irradiated samples, which could be due to photobleaching of the catalyst after extended irradiation times (SI **Figure S4**). Because the Dz-TMR molecule allows for direct fluorescence functionalization of interacting proteins, this technique bypasses the need for traditional Western blotting of biotinylated interactors. Although the fluorescent probe lacks the sensitivity of ECL blot, it is a robust way of comparing catalyst activity between labeled residues along the αS polypeptide chain.

Following these self-labeling tests with αS-C^Ir^_12_, we performed an initial interactome experiment with all five αS-C^Ir^_n_ constructs. The monomeric proteins were incubated in mouse brain lysate (10 µM in 10 mg/mL lysate) and irradiated for 2 minutes in the presence of 250 µM Dz-TMR, then subjected to fluorescence gel analysis. As seen in Figure 3, all five constructs successfully labeled a large number of proteins in the lysates. Significant self-labeling was observed, as seen from the overlay or red TAMRA fluorescence and green Ir catalyst fluorescence (yellow bands). Interestingly, αS-C^Ir^_136_ migrates differently on the gel than the other variants and shows lower levels of self-labeling. From these experiments, constructs αS-C^Ir^_12_, αS-C^Ir^_62_, and αS-C^Ir^_114_ were chosen for further experiments to sample the three regions of αS while providing optimal labeling levels. In particular, among the two NAC regions sites, αS-C^Ir^_62_ outperforms αS-C^Ir^_94_, and among the C-terminal sites, αS-C^Ir^_114_ outperforms αS-C^Ir^_136_, (**Figure 2**).

**Figure 2.**
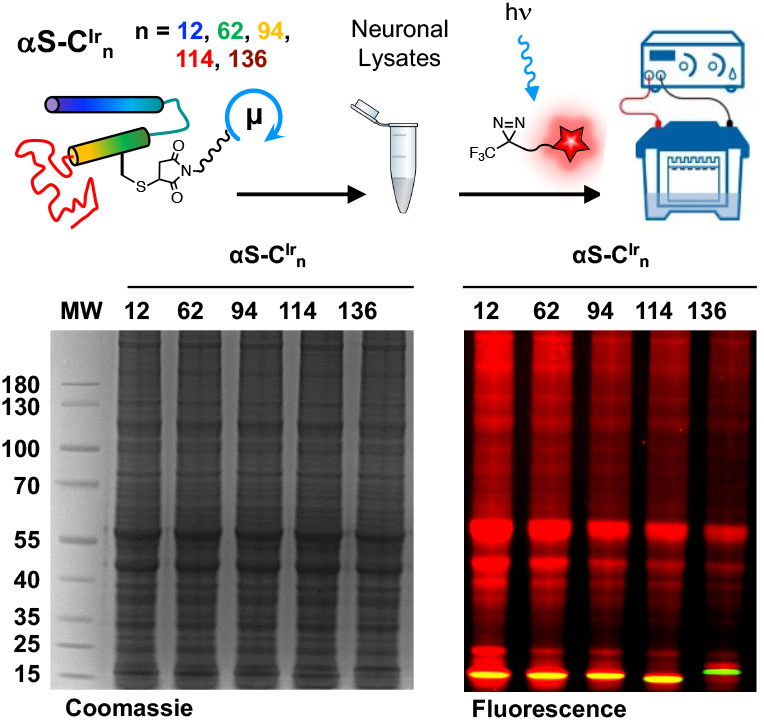
Initial Labeling Tests. Top: Dz-TMR labeling scheme. Bottom: Coommassie-stained and fluorescence gel analysis of αS-C^Ir^ _n_, n = 12, 62, 94, 114, 136 labeling in mouse brain lysates. Arrow indicates αS.

**Figure 3.**
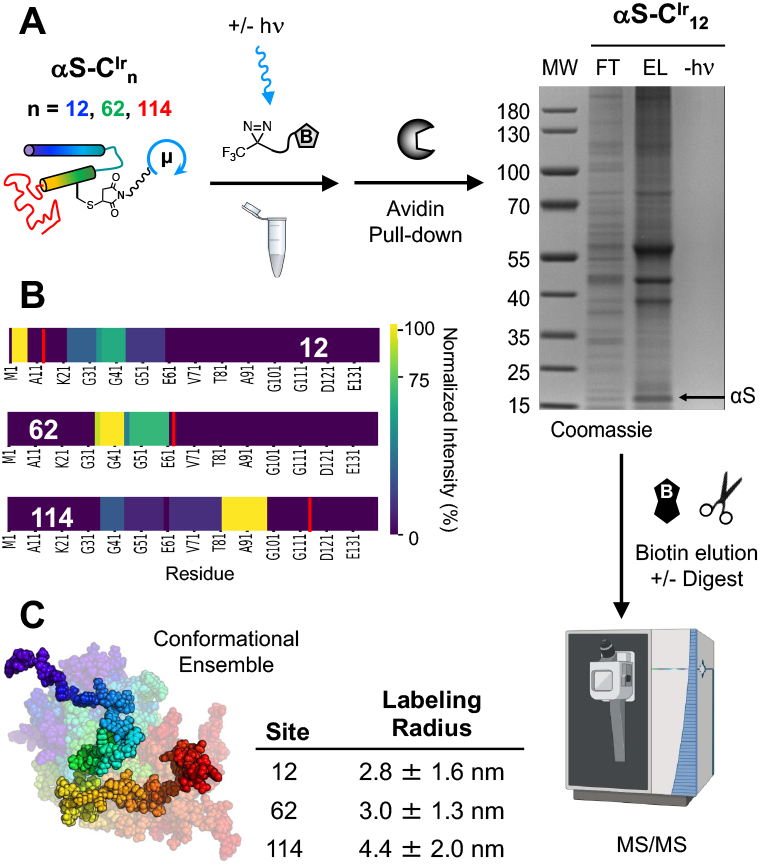
µMap labeling radius determination. (A) µMap workflow using Dz-DTB with monomeric αS-C^Ir^ _n_, n = 12, 62, 114 with gel showing enrichment of specific proteins for αS-C^Ir^_12_ (MW = molecular weight markers, FT = flow through, EL = elution, -hν = unirradiated control), followed by trypsin digestion and LC-MS/MS analysis. (B) Self-labeling by αS-C^Ir^ _n_, n = 12, 62, 114 mapped onto αS sequence with catalyst site indicated by red line and DTB-labelled peptide fragments color-coded according to normalized MS signal intensity. (C) αS monomer conformational ensemble generated in Rosetta for determination of inter-residue distances. Labeling radii determined from average distances of labeled residues to catalyst site for each construct.

Before interactome analysis, we sought to identify labeling events on αS, enabling determination of the µMap labeling radius to inform analysis of interactomes for each labeling site. µMap labeling was conducted for the three chosen αS constructs, incubated at ∼10 µM concentrations (determined based on Ir catalyst absorbance at 316 nm) with 250 µM Dz-DTB in mouse brain lysate. The DTB-labeled proteins were then enriched using streptavidin resin, eluted by incubation with biotin, and digested. In gel analysis of the proximity-labeling reactions, we observed significant enrichment of specific proteins in the elution lanes versus the flow through and virtually no non-specific background in unirradiated controls (αS-C^Ir^_12_ gel in **Figure 3**, αSC^Ir^_62_ and αS-C^Ir^_114_ gels in SI **Figure S5**). For each Ir-labeled variant, all the modified αS peptides were identified using liquid chromatograph tandem MS (LC-MS/MS). The abundances of these peptides were normalized internally, and color-mapped onto the sequence of αS to enable visualization of the labeling radius (**Figure 3**).

From these data, a library of covalently modified amino acid residues was assembled for αS-C^Ir^_12_, αS-C^Ir^_62_, and αS^Ir^-C_114_. Next, a conformational ensemble (n=1000) of αS monomer structures was generated using Fast Floppy Tail^45^ in PyRosetta, and a theoretical distance (Cα-Cα) was calculated from the position of the Ir catalyst to each DTB-labeled residue, for each member of the ensemble. The average radius was computed from the ensemble distance list to approximate the radius of labeling based on experimental data. Theoretical labeling radii can also be obtained easily using literature values for the reactivity of aryl trifluoromethyl diazirines in bulk solvent^46^ (for H_2_O, k = 3.1×10^8^ s^-1^) paired with known diffusion coefficients of similarly sized small molecules (2.0-9.0×10^-6^ cm^2^ s^-1^) and a simple Brownian random walk model.^47^ Comparing these models, the computed radii were remarkably close to the theoretical value for diazirine photo-crosslinking (2-5 nm depending on the diffusion coefficient used). Although our models are still within the range of these theoretical calculations, the slight discrepancy in the αS-C^Ir^_114_ radius could be due to the lack of tryptic cut sites in the C-terminus, biasing statistics to peptides identified farther away from the iridium labeling position. Since αS is an intrinsically disordered protein, this method probably gives a more accurate estimation of the µMap labeling radius than previously demonstrated examples, which were limited by the size and orientation of the protein to which the iridium catalyst was attached.^48^

The absence of the thiopane group in the desthiobiotin compound made it ideal for these competitive pulldowns and radii measurements due to its lower affinity to streptavidin compared to biotin, which allowed us to elute the labeled peptides for labeling site identification by LC-MS/MS. However, subsequent studies using Dz-BTN found it to be superior by providing enhanced sensitivity in chemiluminescent gel assays MS proteomic experiments to identify non-synuclein interactors. In effect, the higher affinity of Dz-BTN for streptavidin allowed for stringent washes to minimize non-specific protein binding to the beads. Additionally, harsh elution conditions from the beads were avoided by trypsin digesting enriched proteins on-bead.

### Catalyst attachment does not perturb αS aggregation

µMap is uniquely well-suited for studies involving small proteins like αS. Traditional APEX/BioID methods require enzyme-protein fusions in which the enzyme conjugate is larger than αS itself –opening the possibility of perturbing its biophysical characteristics. To test the effects of Ir catalyst attachment on fibril formation, recombinant αS-C^Ir^_12_, αS-C^Ir^_62_, and αS-C^Ir^_114_ monomers were aggregated *in vitro* to generate pre-formed fibrils (PFFs). As expected, the Ir-labeled PFFs partitioned into the insoluble pellet after centrifugation and could be visualized by both Coomassie staining and fluorescence gel imaging, with almost no protein remaining in the supernatant (**Figure 4B**). Transmission electron tomography (TEM) images validated the fibrillar nature of the αS-C^Ir^_n_ aggregates, with bulk morphologies similar to wild-type (WT) αS PFFs formed under the same conditions (αS-C^Ir^_12_ PFF TEM in **Figure 4C**, WT αS, αS-C^Ir^_62_, and αS-C^Ir^_114_ PFF TEM in **Figure S6**). These experiments indicate that the chosen Ir catalyst sites are indeed non-perturbing and support their use in interactome studies of the PFFs.

**Figure 4.**
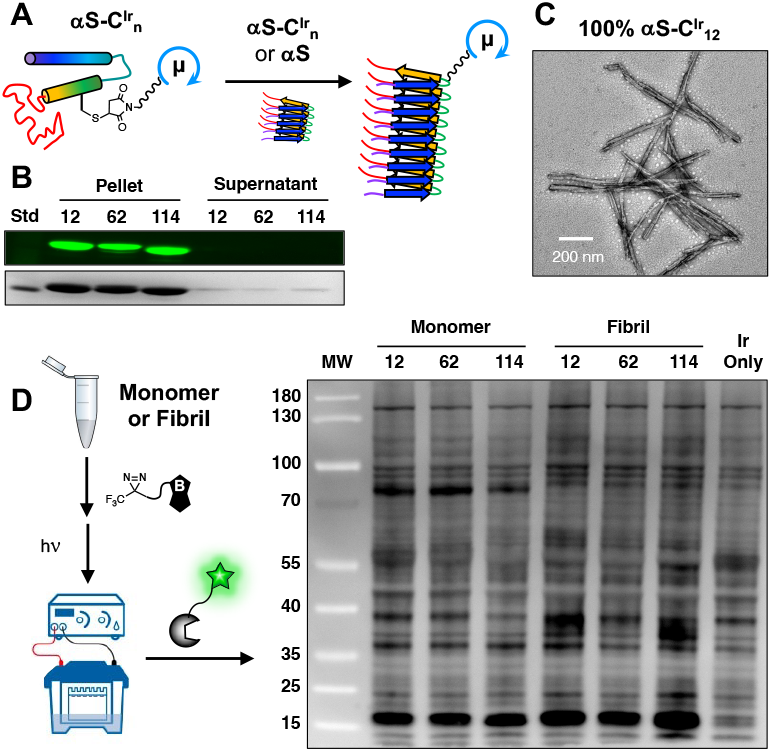
Fibril formation. (A) Aggregation workflow of αS-C^Ir^_n_ where recombinant and labeled monomers were shaken at 350 rpm, 37°C for 72 hours. (B) Iridium fluorescence in pellet fraction over supernatant post-centrifugation for αS-C^Ir^_n_. (C) TEM image of αS-C^Ir^_12_. (D) Labeling workflow for mouse brain lysate enrichment by monomeric and PFF αS-C^Ir^_n_ constructs with Dz-BTN as visualized by STP-HRP chemiluminescence.

Since Ir-labeled αS could still be useful for proximity labeling when the catalyst is only present on a fraction of the proteins in a PFF and would then be even less likely to perturb protein folding and interactions, we also tested fibril formation using mixtures of αS-C^Ir^_n_ proteins with WT αS. TEM analysis of PFFs made with 10% αS-C^Ir^_12_ showed that they also resembled WT αS PFFs and appeared to be homogeneous (SI, **Figure S6**). While this is perhaps not surprising since the 100% αS-C^Ir^_12_ fibrils were unperturbed, it is important to confirm that the αS-C^Ir^_12_ monomers co-fibrillize with the WT αS monomers for any future applications of these 10% labeled PFFs.

We then performed a comparison of proximity labeling in mouse brain lysate using both monomer and 10% Ir-labeled PFFs at 10 µM concentrations of Ir-labeled protein. The mixtures of αS-C^Ir^_n_ and lysate were incubated with 250 µM Dz-BTN and irradiated, then PFF samples were boiled to disaggregate the fibrils and the samples were analyzed in a chemiluminescent gel assay, imaging labeled proteins with a streptavidin horse radish peroxidase conjugate (STP-HRP). As one can see in **Figure 4D**, there are modest differences in the labeling profiles between constructs and between monomer and PFF forms for the same construct. For example, in the monomer lanes, a band at 39 kD is much more prominent in the αS-C^Ir^_12_ and αS-C^Ir^_62_ lanes than the αS-C^Ir^_114_ lane. Additionally, an intense band at 80 kD in the monomer lanes is absent in the PFF lanes. Both monomer and PFF lanes show significant differences with the proteins labeled in the free Ir catalyst control lane. These data show that we can successfully perform proximity labeling with PFFs and that the fibril interactome differs from that of the monomer. However, the large number of labeled proteins in the catalyst lane indicates that MS protein identification experiments must be interpreted with caution and appropriate use of controls to remove false hits.

### Site-specific and conformation-specific αS interactomes

To identify the interactors observed on the gel, αS-C^Ir^_12_, αS-C^Ir^_62_, and αS-C^Ir^_114_ monomers and PFFs were incubated with mouse brain lysate and labeled with a Dz-BTN probe for downstream LC-MS/MS analysis. Using label-free quantification, we compared protein enrichment of the three different labeling positions with each other and with respect to a catalyst-only control, which was added to assess non-specific labeling of abundant proteins in brain lysate. A subtle enrichment in proteins was observed in αS-C^Ir^_12_ samples versus αS-C^Ir^_62_ and αS-C^Ir^_114_. Nonetheless, the bulk proteomic data was consistent across all three positions based on statistical intensity analysis after normalization and imputation (SI, **Figure S7-S10**). Analyses for site-specific and conformation-specific hits were also conducted by sample thresholding, where “specificity” was defined as a protein being detected in *n* replicates of each condition (*n* = 3). From this analysis, we determined 179 monomer-specific hits across all three αS-C^Ir^_12_, αS-C^Ir^_62_, and αS-C^Ir^_114_ positions in direct comparison to only 7 fibril-specific hits (**Figure 5**). This biasing toward monomeric-specific hits is also repeated in position-based comparisons (e.g. αS-C^Ir^_12_ monomer vs. αS-C^Ir^_12_ PFF, **Figure 6**). Interestingly, there are less distinct monomeric protein hits across the three catalyst positions than there are for the same labeled positions for fibrils (**Figure 6**), which can be attributed to possible increased regional selectivity due to the more stable fibril structure.

**Figure 5.**
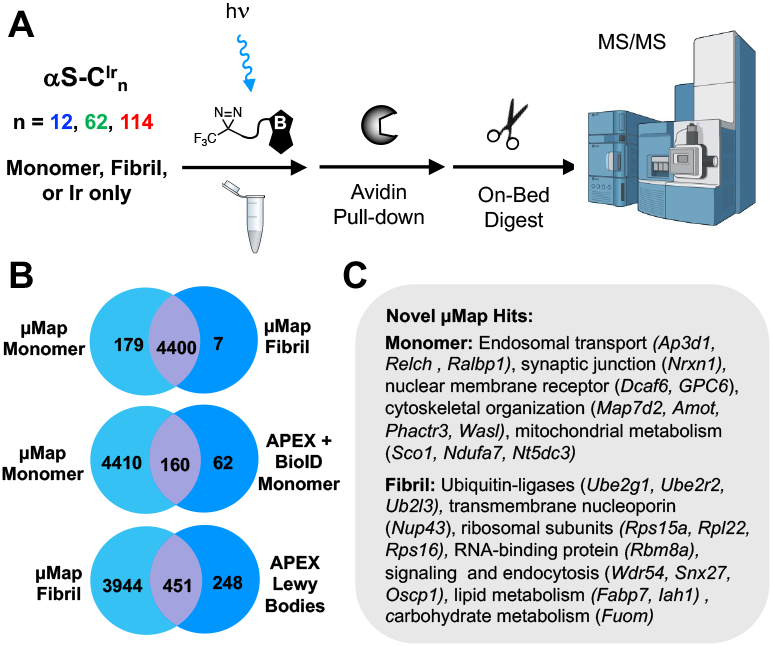
MS/MS analysis of fibril versus monomer enrichment (A) µMap workflow using Dz-BTN with monomeric and fibrillar αS-Ir in lysate. Enrichment by Streptavidin bead pulldown followed by on-bead digest and quantitative MS/MS. (B) Venn diagrams of interactome hits: Top: µMap hits exclusive to pooled samples across all monomeric positions (All Monomer) vs pooled αS-Ir, fibril samples (All Fibril). Middle: Venn diagram of monomeric interactome hits found by µMap as compared to previously described monomeric hits found by APEX in rat cortical neurons.^37^ Bottom: Venn diagram of PFF interactome hits found by µMap as compared to hits previously described from LB in human brain tissue.^38^ (C) Novel µMap proteins and their cellular pathways not previously described by either monomeric or LB experiments.

**Figure 6.**
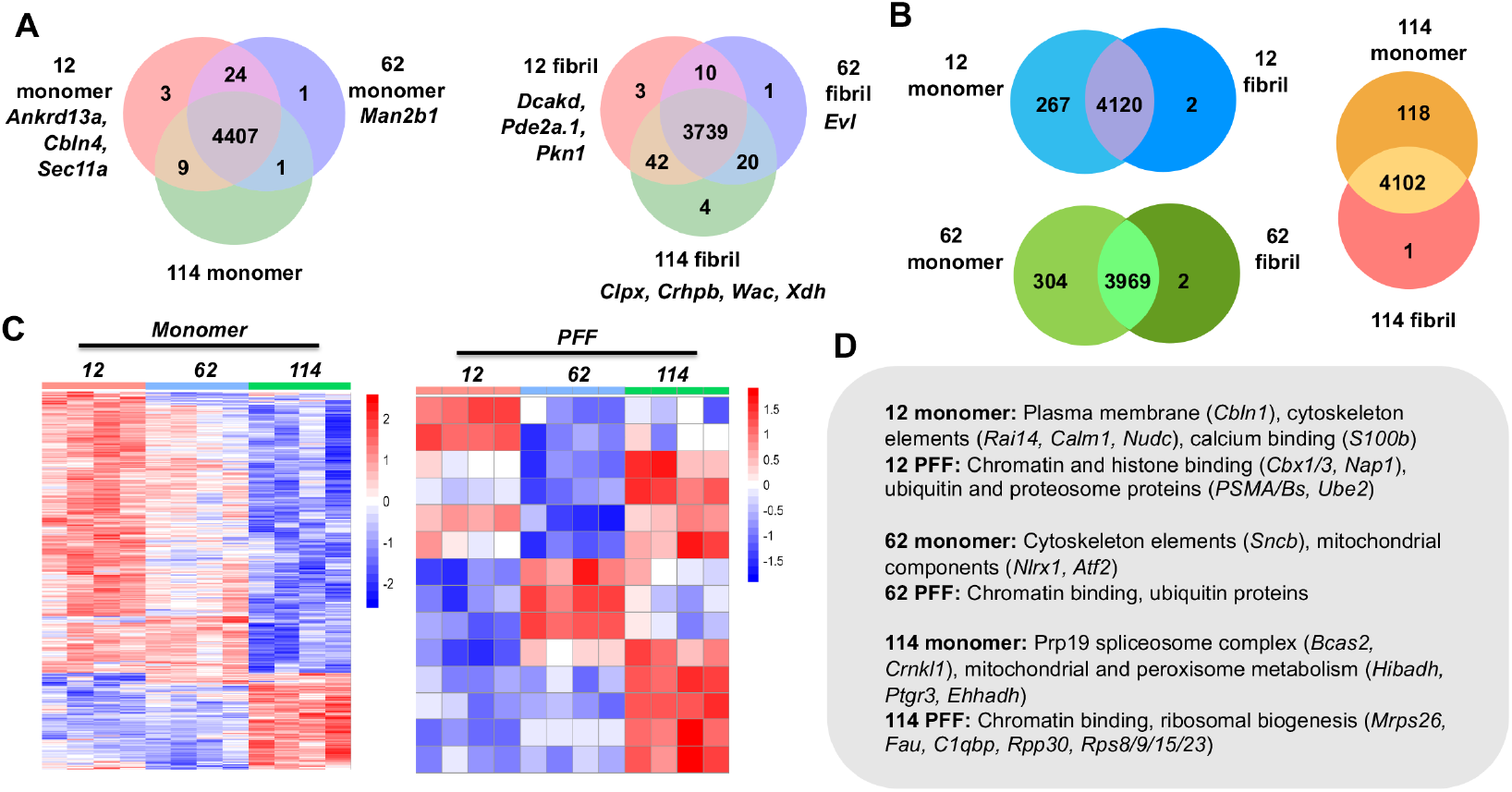
Site and Structure Specific Interactome Comparisons. (A) Site-specific comparisons between αS-C^Ir^_12_, αS-C^Ir^_62_, and αS-C^Ir^_114_ monomers (left) and PFFs (right). (B) Venn diagram of conformation-specific hits (monomer vs. PFF) by labeling site. (C) Heatmap of relative enrichment across all sites for monomers and PFFs. Scaled counts normalized on log2 scale with relative enrichment as blue and vice versa as red. (D) Representative list of enriched hits by site, conformation, and cellular component.

For monomer interactomes, we identified proteins involved in endosomal transport processes, cytoskeletal organization, and mitochondrial metabolism across all monomeric constructs. These cellular networks have been previously implicated in αS trafficking and intracellular fate.^49-52^ Key protein differences were also observed across all samples when compared to the enriched proteins in the Ir-only control. A prominent hit enriched for both the N-terminal and NAC domain constructs (αS-C^Ir^_12_/αS-C^Ir^_62_) was S100β, a calcium-binding protein that exists as a homodimer and is primarily found in the astrocytes (**Figure 6**, SI **Figures S11, S13**). Previous studies have implicated high concentrations of S100β as a biomarker for a myriad of different neurodegenerative diseases. Inhibition of astrocytic production of S100β has been shown to improve amyloid-β deposits with implication in AD.^53^ While S100β interactions with αS have not been shown to date, S100β has been shown to be upregulated in PD brain tissue, potentially modulating dopaminergic neuron function and neuroinflammation.^54^ The highly positively charged N-terminus, measured by the high isoelectric (pI) point of residues 1-60 of αS (pI = 9.52), may facilitate preferential interaction to S100β, with the low isoelectric point 4.66. However, prior research suggests that S100β contributes to dysregulation of extracellular signaling mechanism with little evidence that the protein exists in measurable amounts intracellularly in dopaminergic neurons. Thus, further investigation must be done to validate the mechanism of how these proteins may affect neurobiology and/or neurodegeneration. From Gene Ontology (GO) analysis, both the N-terminal and NAC regions have been shown to label proteins involved in the microtubule cytoskeleton over promiscuous labeling from the Ir-only control (SI, **Figures S12, S14**). αS-C^Ir^_12_ and αS-C^Ir^_62_ monomer also showed the greatest amount of shared hits, further supporting the overlapping radii between the two labeled regions (**Figure 6**). In terms of C-terminal hits, glutamate dehydrogenase 1 (Glud1) shows an enrichment in αS-C^Ir^_114_ samples against both N-terminal and NAC domain mutants (SI, **Figure S16**). Glud1 is a mitochondrial enzyme that catalyzes the conversion of glutamate to α-ketoglutarate, but is novel in αS-interactome studies. This protein has broad implications in neurodegenerative disease due to its role in mitochondrial metabolism, as its dysregulation has been shown to be correlated with numerous disorders. Interestingly, GO analysis of αS-C^Ir^_12_ and αS-C^Ir^_114_ (SI, **Figures S14, S16**), two monomeric regions that are spatially further apart, also show a shared enrichment for spliceosomal proteins, although the biological significance of these proteins in relation to αS must be validated in both cellular and animal models rather than in whole brain lysate.

Analysis of protein enrichment from αS PFF pulldowns samples also suggests the existence of conformation-specific αS interactors. Pairwise fold-change comparisons between αS-C^Ir^_12_, αS-C^Ir^ _62_, and αS-C^Ir^ _144_ PFFs with Ir-only control demonstrate that the bulk proteomic labeling of αS Ir PFF in comparison to promiscuous catalyst labeling is significantly lower (SI, **Figures S16, S19, S21**). Although there is a myriad of physical explanations for this phenomenon (limitations due to fibril light scattering, conformation of the catalyst in the fibril cleft, probe access), it is also important to consider that the Ir-catalyst only control is a measure of abundant and non-specific interactors, and it is therefore reasonable to consider proteins enriched in PFF αS-Ir as fibril-selective. In addition to selectivity, more enriched hits seem to be shared overall between the labeled regions in the fibrils versus the monomeric constructs, which could suggest morphology-specific interactions (**Figure 6**).

### Comparison to previous interactome studies

We then compared our µMap PL proteomic dataset to previous APEX-based PL studies of αS monomer conducted in rat cortical neurons (**Figure 5B**). APEX hits were BLAST-sequence aligned to find the top scoring mouse homologues and cross-referenced to the µMap-enriched proteome. As expected, a large majority of APEX hits were found to be shared by our µMap dataset. Within these comparisons, we were additionally able to identify novel hits related to the enriched cellular networks. One such protein, Neurexin-1, is a synaptic protein implicated in the neuron-to-neuron αS propagation.^55^ Further studies using methods like µMap that afford tight spatiotemporal control could elucidate the exact mechanisms of αS internalization.

PFFs were initially identified as a transitionary structure between monomers and pathological LBs. As such, it is possible that probing PFF interactomes could give novel insight into the molecular mechanisms of disease progression and propagation. With this in mind, we compared LB APEX PL hits derived from PD and Lewy Body Disease (LBD) brain tissue to our αS-Ir PFF µMap interactomes. We verified that the majority the LB APEX hits were also found in our analysis. In addition, we were also able to identify novel hits in ribosomal subunits and RNA-binding proteins. Furthermore, we identified novel proteins involved in both short and long chain fatty acid hydrolysis (Fabp7, Iah1) as well as the fucose metabolic protein Fuom, suggesting αS’s role in lipid and carbohydrate metabolism and subsequent dysregulation in disease states (**Figure 5**).^56-57^ GO analysis of only terms enriched by αS-Ir PFFs enriches for both proteosomal complex proteins and, intriguingly, DNA-binding proteins across all three labeling sites (SI, **Figures S17, S18, S10, S22**). αS has been previously implicated in a variety of epigenetic roles, and these interactions may be involved.^58^ In addition to this finding, the top enriched protein hit in all αS-C^Ir^_12_, αS-C^Ir^_62_, and αS-C^Ir^_114_ PFFs conditions was Nup43, a protein subcomponent of the membrane-embedded nucleoporin subcomplex Nup107-160 (SI, **Figure S23**). While no such interactions with Nup43 have yet been found, αS has previously been suggested to localize into the nucleus and bind to DNA, with one model suggesting that the aggregation of cytoplasmic synuclein decreases the amount in nuclear loci.^59^ These results are especially interesting compounded with the enrichment of proteosomal proteins in PFF samples; in particular, αS-C^Ir^_12_ PFF highly enriches for PSMA4, a subunit protein in the 26S proteosome (**Figure 6**). In support of our proteomic findings, the N-terminal region is essential for peptic cleavage by the proteosome.^60^ The inhibition of the proteosomal complex has been shown to decrease the clearance of intracellular αS aggregates seeded by recombinant PFFs.^61^ One study suggests that αS is an inhibitory substrate, in which its binding to the proteosome and/or sequestering of other proteosomal substrates leads to a positive feedback mechanism of increased cytoplasmic αS aggregates.^62^

### Hit validation

While an exhaustive validation of every hit from our monomer and PFF interactomes is not feasible in this initial report, we wished to demonstrate several mechanisms for further analysis from the hits discussed above. First, for Western blotting, mouse brain lysate (5µg/µL) was irradiated with 10 µM αS and 500µM Dz-BTN for αS-Ir monomer and PFF constructs. Labeled lysate proteins were subsequently enriched using streptavidin magnetic beads, run on a gel, and transferred onto a Western blot for proteomic hit validation. As a proof-of-concept validation of the previous proteomics data, the calcium-dependent S100β was enriched by αS-Ir monomer, but not αS-Ir fibril, in comparison to WT and the positive lysate-only control (SI, **Figure S24**). Other hits such as Glud1 and JCAD1were also analyzed in the same fashion (SI, **Figure S24**). As a second validation method, super-resolution imaging by Dual-Color DNA Points Accumulation for Imaging in Nanoscale Technology (DNA-PAINT) was used to visualize and quantify localization overlap of Glud1 and αS in hippocampal neurons. Mouse hippocampal neurons were used as a cell system to measure endogenous αS-Glud1 interactions. Standard cell culture protocol was followed by DNA-PAINT immunostaining (SI, **Table S1**). Quantitative analysis via a custom script was used calculate the percentage of localization overlap of αS onto Glud1. Glud1 signal was primarily localized to mitochondrial substructures and had significant overlap with endogenous levels of αS in which 18.2% of all mitochondrial structures labeled with Glud1 were found to be enriched with αS (SI, **Figure S25**). Thus, these orthogonal techniques validate novel hits observed in our µMap interactomes, including the specificity of the interactions between monomer and PFF.

### µMap interactome labeling in hippocampal neurons

Primary neurons have previously been shown to uptake exogenous fluorescently-labeled αS.^63-64^ Similarly, we performed proof-of-concept µMap interactomics experiments *in cellulo* as shown in **Figure 7**. These studies demonstrated both successful intercellular uptake of Ir-labeled αS and photo-induced labeling via Dz-BTN in all labeled sites for both monomers and PFFs (**Figure 7** and SI, **Figure S26**). Neurons that were treated without αS-Ir (WT or no αS treatment) or Dz-BTN probe did not show significant photolabeling. The interactome of αS PFFs was seen to be distinctly puncta-like with colocalization to the early endosomal marker LAMP1, consistent with other work implicating the endosomal pathway in αS PFF internalization.^64^ Fluorescent signal also was dependent on the concentration of Dz-BTN probe more so than the amount of αS-Ir, consistent with its catalytic role (SI, **Figures S27, S28**). These results suggest the compatibility of µMap labeling within the complexity of the intracellular environment and act as a proof-of-concept towards future µMap PL studies in physiological neuron environment.

**Figure 7.**
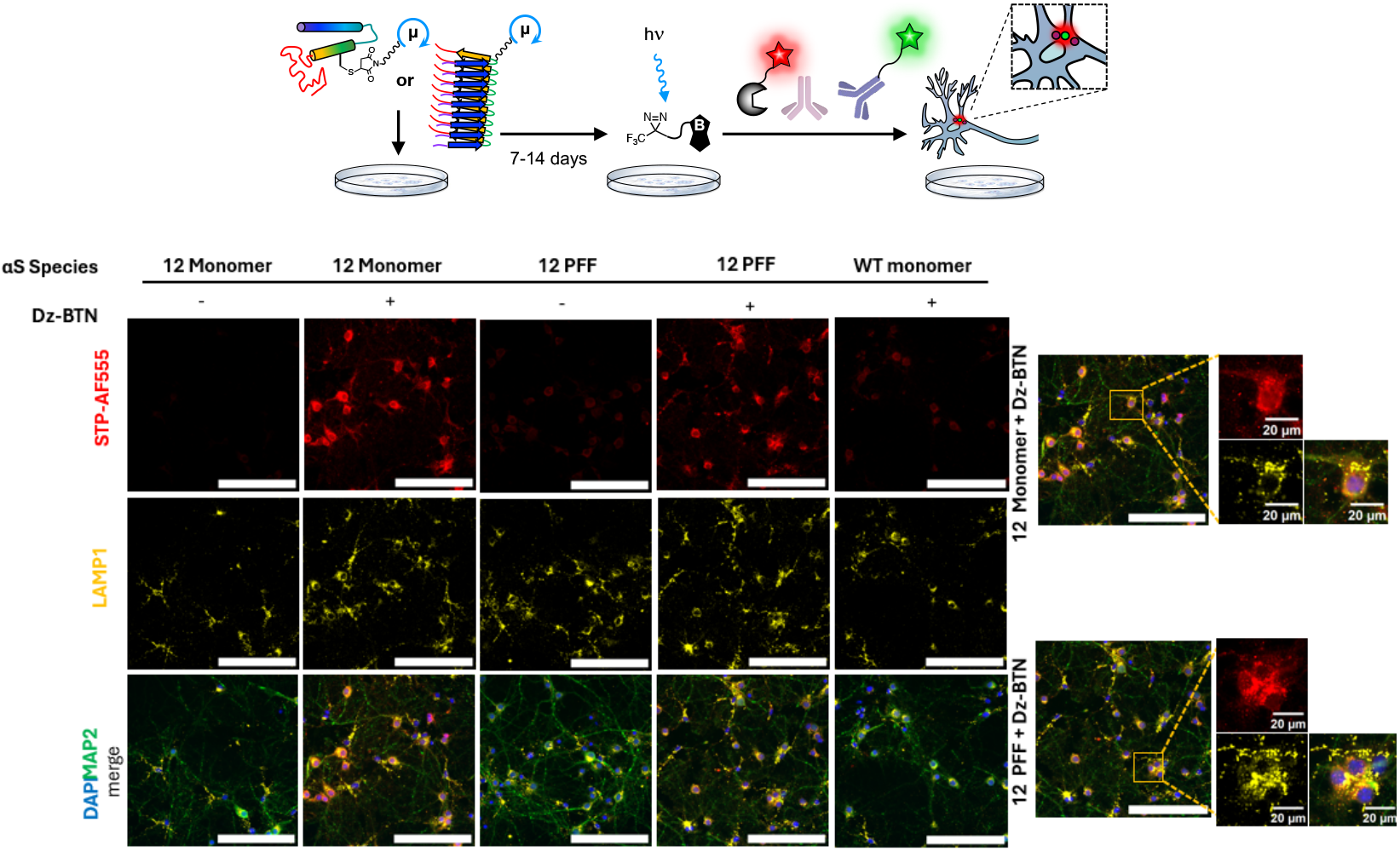
Top: µMap interactome labeling in primary hippocampal neurons. (A) Treatment of neonate hippocampal neurons with αS-Ir followed by Dz-BTN photo-crosslinking after incubation. Tagged neuronal proteins are visualized by fluorescent immunocytochemistry. Bottom: Biotinylated αS-Ir interactome shown using fluorescent STP-AF555 (red). Interactome colocalization visualized with endosomal marker LAMP1 (yellow). Neuron morphology shown by MAP2 (green) and DAPI (blue). All scale bars 150 µm unless otherwise indicated.

## Conclusion

µMap is unique amongst PL methods in its tight labeling radius and small (∼1 kDa) biorthogonally incorporable PL motif, which allows for site-specific conjugation. However, µMap is still a new method, and relatively few labs have yet adopted it. We sought to integrate the µMap platform in our own PL studies and demonstrate the ease with which this transformative technology can be incorporated into experiments in chemical biology. To highlight this, we developed a cysteine-reactive version of the µMap iridium catalyst and optimized labeling on our model protein αS. Novel diazirine probe partners for µMap were synthesized, and these have implications in improved proteomic workflows (Dz-DTB) and fluorescence labeling of substrate and interactor proteins (Dz-TMR). Despite a tight theoretical labeling radius, experimental validation often has been limited by orientation of the antibody-µMap platform (exp. radius = 50 nm, theo. radius = 4 nm). A hybrid computational and experimental approach was utilized that enabled determination of µMap labeling radii for various αS-Ir labeled constructs, highlighting the unprecedented tight radius of this PL method. By incorporating µMap directly on αS and using our Dz-DTB probe, we were able to experimentally obtain a labeling radius remarkably close to the theoretical value (2-4 nm). Next, by incorporating the catalyst at different positions on αS monomer and fibril, we identified proteins that preferentially interact with the three different regions of both PFF and monomeric αS in a total mouse brain homogenate. Protein-protein interactions were identified whose novelty relative to previous αS PL experiments are likely a direct consequence of the site-specific incorporation and small size of the µMap photocatalyst. We have performed a set of exemplary experiments to demonstrate how the interactome hits can be pursued in more depth and will continue to pursue additional hits in this fashion for subsequent reports. Finally, we were able to demonstrate a robust labeling *in cellulo* using primary hippocampal neurons. These studies serve as an additional proof-of-principle of this system and help to show how it can be incorporated into chemical biology workflows in the future.

## ACKNOWLEDGMENT

This research was supported by the National Institutes of Health (NIH RF1-NS103873 to E.J.P.). Instruments supported by the NIH and NSF include NMR (NSF CHE-1827457), mass spectrometers (NIH S10-OD030460), and a computing cluster (NIH S10-OD023592). M.G.L. was supported by an Age Related Neurodegenerative Disease Training Grant fellowship (NIH T32AG000255). E.Y. was supported by the NIH Chemistry Biology Interface Training Program (T32-GM133398). Z.L. thanks a Washington University School of Medicine BMB Seed Grant (PJ000027587) and support from the Research Education Component through the P30-AG066444 grant.

