## Supporting Information for "Determination of α-Synuclein Protein Interactions by μMap Photo-proximity Labeling"

### General

**Materials.** *E. coli* BL21(DE3) cells were purchased from Stratagene (La Jolla, CA, USA). Milli-Q filtered (18 MΩ) water was used for all solutions (Millipore; Billerica, MA, USA). Cytiva Vivaspin centrifugal filter units (3 kDa MWCO) were purchased from Cytiva Life Sciences (Marlborough, MA, USA). All other reagents and solvents were purchased from Fisher Scientific (Pittsburgh, PA, USA) or Sigma-Aldrich unless otherwise specified. DNA sequencing was performed at the University of Pennsylvania DNA sequencing facility. Synthetic reactions were monitored by thin-layer chromatography (TLC) on pre-coated silica 60 F254 aluminum plates (MilliporeSigma, Burlington, MA, USA), and spots were visualized by ultraviolet (UV) light.

**Instruments.** Evaporation of solvents was performed under reduced pressure at 40 °C using a rotary evaporator. Flash column chromatography was performed on a Biotage® (Charlotte, NC, USA) Isolera One system equipped with Biotage® SNAP KP-Sil cartridges. Nuclear magnetic resonance (NMR) spectroscopy was performed on a Bruker (Billerica, MA, USA) Avance Neo 600 (600 MHz for $^{1}$H and 150 MHz for $^{13}$C) with chemical shifts (δ) reported in parts per million (ppm) relative to the solvent ($CDCl_3$, $^{1}$H 7.26 ppm, $^{13}$C 77.16 ppm; dimethyl sulfoxide (DMSO)-$d_6$, $^{1}$H 2.50 ppm, $^{13}$C 39.52 ppm). Low resolution liquid chromatography mass spectrometry (LCMS) was carried out using a Waters (Milford, MA, USA) SQD equipped with an Acquity UPLC instrument in positive-ion mode. High resolution mass spectrometry (HRMS) for small molecules was obtained on a Waters LCT Premier XE LC/MS system. Matrix-assisted laser desorption/ionization with time-of-flight detector (MALDI) mass spectra were acquired on a Bruker rapifleX LRF instrument (Billerica, MA, USA). Fluorescence spectra and protein quantification measurements were collected with a Tecan SPARK plate reader (Mannedorf, Switzerland). Initial protein purification performed on Cytiva Life Sciences ÄKta FPLC system (Marlborough, MA, USA). Final protein purification performed on an Agilent 1260 Infinity II HPLC (Santa Clara, CA, USA) on a Phenomenex C4 Reverse Phase Semi-prep column (Torrance, CA, USA).

### Small Molecule Synthesis and Characterization

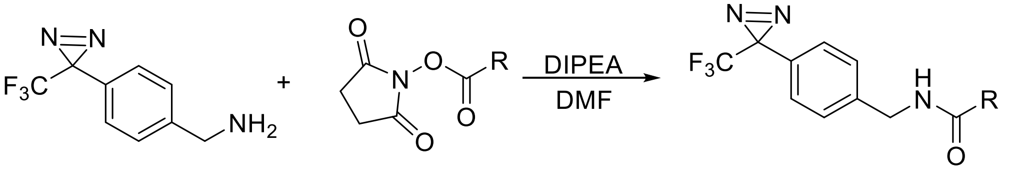


**General Procedure A. (for simple preparation of diazirine probe molecules from commercially available succinimidyl esters)** (4-(3-(trifluoromethyl)-3*H*-diazirin-3-yl)phenyl)methanamine (1.0 eq.) was added to a solution of the appropriate succinimidyl ester (1.0 eq.) in DMF and DIPEA (2.0 eq.). The reaction was allowed to stir for 10-12 hours. The resulting reaction mixture was concentrated under reduced pressure and purified directly using Biotage normal phase silica chromatography (0-20% MeOH in DCM).

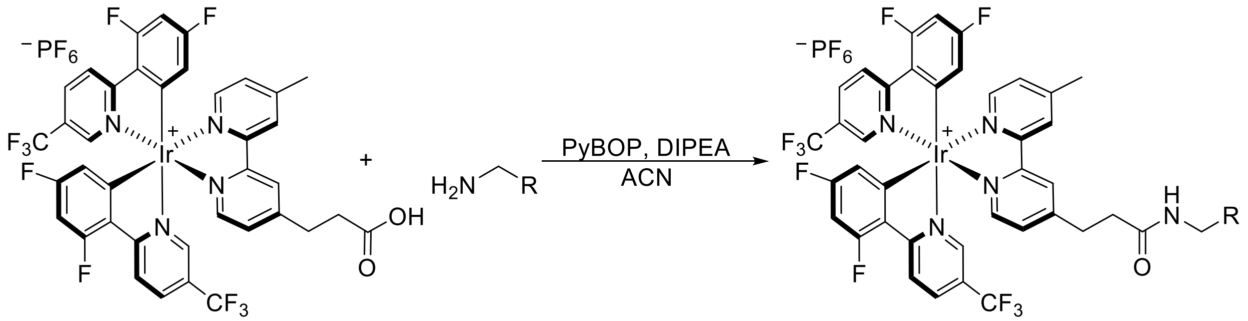


**General Procedure B. (for preparation of iridium catalysts)** A 4 ml scintillation vial was flame-dried under vacuum with a stir bar. After allowing the reaction vessel to cool, 100 mg (0.0912 mmol, 1.0 eq.) of **Ir-COOH**, prepared as described previously[1], was added to the vial followed by 95 mg of benzotriazole-1-yloxytripyrrolidinophosphonium hexafluorophosphate (PyBOP) (0.1825 mmol, 2.0 eq.), the desired biorthogonal handle as a primary amine (0.1825 mmol, 2.0 eq.), 100 µl of diisopropylethylamine (DIPEA) (0.574 mmol, 6.3 eq.), and 1.0 ml of ACN. The reaction was allowed to stir overnight at room temperature. The crude reaction was transferred to a 100 ml separatory funnel and ~30 ml of ethyl acetate (EtOAc) was added. The organic layer was washed twice with saturated $NaHCO_3$, twice with 0.1 M HCl, and twice with saturated NaCl. The resulting organic layer was dried using $Na_2SO_4$ and concentrated under reduced pressure. The resulting solid was re-suspended in cold diethyl ether ($Et_2O$) and collected via filtration through a Buchner funnel. The solid residue was washed three times with cold $Et_2O$, collected in a scintillation vial, and dried under high vacuum to yield the pure product as a yellow solid.

**1-(5-((3aR,4R,6aS)-2-oxohexahydro-1H-thieno[3,4-d]imidazol-4-yl)pentanamido)-N-(4-(3-(trifluoromethyl)-3H-diazirin-3-yl)benzyl)-3,6,9,12-tetraoxapentadecan-15-amide (Dz-BTN)**

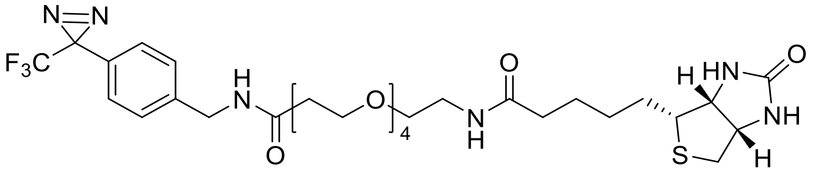

**Dz-BTN** was synthesized using **general procedure A**, following a published procedure[2] in 94% yield from biotin-PEG4-succinimidyl ester. **$^1$H NMR (600 MHz, DMSO)** δ 8.46 (t, *J* = 6.0 Hz, 1H), 7.85 (t, *J* = 5.7 Hz, 1H), 7.38 (d, *J* = 8.5 Hz, 2H), 7.23 (d, *J* = 8.1 Hz, 2H), 6.42 (s, 1H), 6.37 (s, 1H), 4.30 (m, *J* = 6.0 Hz, 3H), 4.12 (m, *J* = 7.9, 4.5, 1.9 Hz, 1H), 3.62 (t, *J* = 6.3 Hz, 2H), 3.49 (d, *J* = 3.0 Hz, 12H), 3.38 (t, *J* = 6.0 Hz, 2H), 3.18 (m, *J* = 5.6 Hz, 2H), 3.09 (m, *J* = 8.6, 6.2, 4.4 Hz, 1H), 2.81 (dd, *J* = 12.4, 5.1 Hz, 1H), 2.58 (d, *J* = 12.4 Hz, 1H), 2.38 (t, *J* = 6.3 Hz, 2H), 2.06 (t, *J* = 7.5 Hz, 2H), 1.61 (ddt, *J* = 13.5, 9.8, 6.1 Hz, 1H), 1.54 – 1.41 (m, 3H), 1.36 – 1.24 (m, 4H). **$^{13}$C NMR (151 MHz, DMSO)** δ 172.60, 170.81, 163.20, 142.73, 128.48, 126.87, 126.30, 123.31, 121.49, 70.25, 70.19, 70.02, 69.63, 67.30, 61.51, 59.68, 55.89, 53.75, 49.05, 42.00, 40.31, 38.90, 36.59, 35.56, 28.66, 28.50, 25.73.

**6-(5-methyl-2-oxoimidazolidin-4-yl)-N-(4-(3-(trifluoromethyl)-3H-diazirin-3-yl)benzyl)hexanamide (Dz-DTB)**

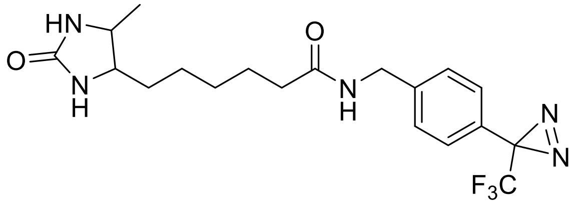

**Dz-DTB** was synthesized from 2,5-dioxopyrrolidin-1-yl 6-((4R,5S)-5-methyl-2-oxoimidazolidin-4-yl)hexanoate using **general procedure A**. The product was isolated as a pure white powder in 98% yield. **$^1$H NMR (600 MHz, $CDCl_3$)** δ 7.30 (d, *J* = 8.2 Hz, 2H), 7.12 (d, *J* = 8.0 Hz, 2H), 6.87 (t, *J* = 6.0 Hz, 1H), 6.11 (s, 1H), 4.77 (s, 1H), 4.47 – 4.31 (m, 2H), 3.75 (p, *J* = 6.7 Hz, 1H), 3.64 – 3.51 (m, 1H), 2.22 (t, *J* = 7.2 Hz, 2H), 2.11 (d, *J* = 7.2 Hz, 1H), 1.66 (p, *J* = 7.0 Hz, 2H), 1.43 – 1.19 (m, 7H), 1.05 (d, *J* = 6.5 Hz, 3H). **$^{13}$C NMR (151 MHz, $CDCl_3$)** δ 173.17, 163.98, 140.79, 128.19, 128.05, 126.70, 123.01, 121.19, 55.90, 51.36, 42.78, 35.54, 29.46, 28.16, 25.51, 25.16, 15.69. **LCMS** calculated for $[M+H]^+$ = 412.195 observed = 412.323.

***N*-(9-(2-carboxy-4-((4-(3-(trifluoromethyl)-3*H*-diazirin-3-yl)benzyl)carbamoyl)phenyl)-6-(dimethylamino)-3*H*-xanthen-3-ylidene)-N-methylmethanaminium (Dz-TAMRA)**

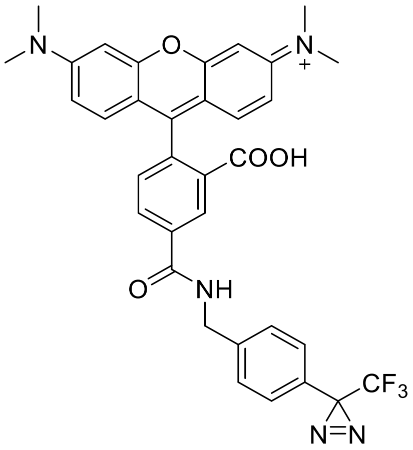


**Dz-TAMRA** was synthesized from TAMRA succinimidyl ester using **general procedure A**. The product was isolated as a pink solid. $^{1}$**H NMR** (600 MHz, DMSO) δ 9.51 (t, *J* = 5.9 Hz, 1H), 8.51 (s, 1H), 8.28 (dd, *J* = 8.0, 1.6 Hz, 1H), 7.51 (d, *J* = 8.1 Hz, 2H), 7.31 (dd, *J* = 33.2, 8.0 Hz, 3H), 6.52 (p, *J* = 8.9 Hz, 6H), 4.56 (d, *J* = 5.8 Hz, 2H), 2.95 (s, 12H). $^{13}$**C NMR** (151 MHz, DMSO) δ 168.25, 164.80, 152.28, 152.12, 141.99, 135.68, 134.44, 128.49, 128.32, 127.07, 126.54, 126.03, 124.46, 123.49, 122.85, 121.03, 109.16, 105.71, 97.92, 54.92, 42.45, 28.18, 27.92. **LCMS** calculated for $[M]^+$ = 628.217 observed = 628.668.

**Ir-COOH**

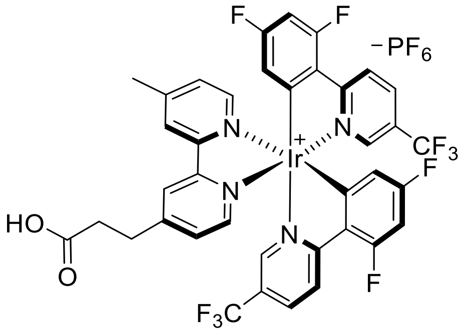


**Ir-COOH** was prepared as described previously.[1] Briefly, a flask was charged with Ir[dF(CF$_3$)ppy]MeCN$_2$•PF$_6$ and 3-(4'-methyl-[2,2'-bipyridin]-4-yl)propanoic acid. Dichloromethane (DCM) and methanol (MeOH) (4:1) were added and the reaction was stirred at 30 °C for 16 hours. The reaction was concentrated under reduced pressure and purified by normal phase silica column chromatography (0-5% MeOH in DCM). (68% yield, yellow solid) $^{1}$**H NMR** (600 MHz, DMSO) δ 12.35 (s, 1H), 8.83 (dd, *J* = 4.0, 1.7 Hz, 2H), 8.45 (tt, *J* = 11.1, 8.9 Hz, 4H), 7.82 (dd, *J* = 11.7, 5.7 Hz, 2H), 7.67 (dd, *J* = 2.2, 1.1 Hz, 1H), 7.64 (dd, *J* = 5.8, 1.7 Hz, 1H), 7.58 (dd, *J* = 5.7, 1.7 Hz, 1H), 7.50 (dd, *J* = 2.1, 1.1 Hz, 1H), 7.07 (ddd, *J* = 12.2, 9.4, 2.4 Hz, 2H), 5.77 (ddd, *J* = 9.7, 8.2, 2.4 Hz, 2H), 3.06 (t, *J* = 7.5 Hz, 2H), 2.74 (t, *J* = 7.5 Hz, 2H), 2.57 (s, 3H). $^{13}$**C NMR** (151 MHz, DMSO) δ 173.08, 166.78, 164.66, 163.03, 162.95, 155.57, 155.31, 155.13,

154.94, 152.60, 150.24, 149.95, 137.57, 129.66, 128.93, 126.44, 125.96, 125.09, 124.64, 123.75, 123.61, 122.84, 122.78, 121.03, 114.18, 114.06, 99.78, 99.60, 99.42, 33.12, 29.80, 20.94. **LCMS** calculated for $[M]^+$ = 949.134 observed = 949.814.

**Ir-Mal**

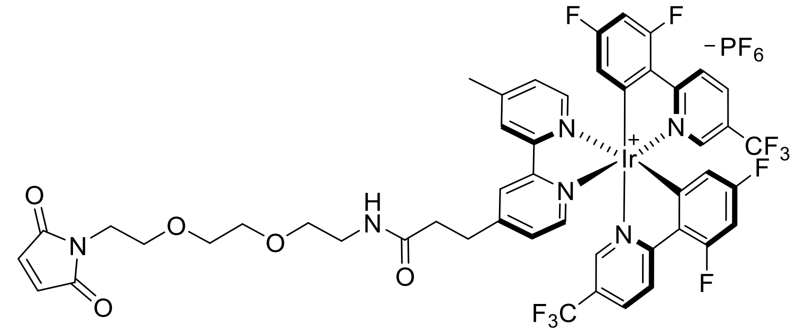


**Ir-Mal** was synthesized from $H_2N$-PEG2-Maleimide using **general procedure B**. Product was isolated as a yellow solid in 55% yield. **$^1$H NMR** (600 MHz, $CDCl_3$) δ 8.70 (t, *J* = 11.5 Hz, 2H), 8.48 (ddd, *J* = 17.6, 8.9, 3.3 Hz, 2H), 8.05 (dd, *J* = 14.9, 8.6 Hz, 2H), 7.75 (dd, *J* = 13.2, 5.3 Hz, 2H), 7.60 (d, *J* = 14.8 Hz, 1H), 7.55 – 7.41 (m, 2H), 7.35 (d, *J* = 5.8 Hz, 1H), 6.70 (s, 1H), 6.68 – 6.58 (m, 3H), 5.69 – 5.55 (m, 2H), 3.71 (t, *J* = 5.6 Hz, 2H), 3.64 (t, *J* = 5.6 Hz, 2H), 3.59 (d, *J* = 4.5 Hz, 2H), 3.54 (d, *J* = 4.6 Hz, 2H), 3.48 (t, *J* = 5.5 Hz, 2H), 3.36 (d, *J* = 5.5 Hz, 2H), 3.21 (d, *J* = 6.5 Hz, 2H), 3.01 – 2.88 (m, 1H), 2.81 (qd, *J* = 15.4, 7.4 Hz, 2H), 2.67 (s, 3H).

**Ir-DBCO**

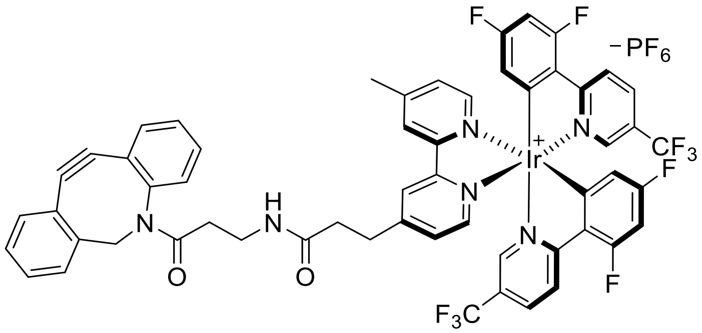


**Ir-DBCO** was synthesized as described previously, and using **general procedure B**.[2] This compound was synthesized and characterized by Vinayak Pagar.

### αS Expression, Purification, and Labeling

**αS Cysteine and AzF Mutant Expression.** α-Synuclein (αS) was expressed, purified, and aggregated as previously described, with slight modifications.[3] For cysteine mutant constructs, human αS with a site-specific cysteine mutant and C-terminal intein-$His_6$ fusion was transformed into *Escherichia coli* (*E. coli*) BL21 cells and plated on ampicillin (amp) plates (100 µg/ml). For AzF mutants, αS with the amber codon (TAG) at the desired position was transformed into BL21 (DE3) cells engineered with a pDULE2-*p*XF (*p*-X-phenylalanine) plasmid containing the *M. jannaschii* mutant *p*-XF-tRNA synthetase and $tRNA_{CUA}$ and was plated on plates supplemented with amp (100 µg/ml) and streptomycin (strep) (100 µg/ml). A single colony was then inoculated into a 5 ml primary culture containing amp (cysteine mutants) or amp/strep (AzF mutants) (100 µg/ml) in Luria-Bertain (LB) media and grown for 5-6 h with shaking (250 rpm) at 37 °C. The primary culture was then transferred to 1 L LB containing amp (cysteine mutants) or amp/strep (AzF mutants) (100 µg/ml) and grown as previously described until reaching an optical density ($OD_{600}$) of 0.8-1.0. At this stage, protein production was induced by the addition of isopropyl β-D-1-thiogalactopyranoside (IPTG) to 1 mM final concentration, and the culture was grown with shaking (250 rpm) overnight at 18 °C. The following day, cells were harvested by centrifugation at 4 °C for 20 min at 4000 rpm (Sorvall GS3 rotor). Pellets were then re-suspended in 20 ml/L culture of 40 mM Tris, pH 8.3 supplemented with EDTA-free protease inhibitor tablets (Pierce Biotechnology; Waltham, MA, USA) and transferred to a metal cup for sonication. Cells were lysed by sonication on ice with a Q700 probe sonicator (QSonica LLC; Newtown, CT, USA) with the following settings: Amplitude 50, Process Time 2-3 min, Pulse-ON Time 1 s, Pulse-OFF Time 1 s. Crude lysate was then transferred to 50 ml centrifugation tubes and clarified via centrifugation at 14,000 rpm for 45 min (Sorvall SS34 rotor). Following centrifugation, supernatant was removed and transferred to a 50 ml Falcon tube. 5 ml of nickel agarose resin (GoldBio; St. Louis, MO, USA) was added, and the lysate-nickel mixture was incubated with nutation at 4 °C for 1-2 h. Lysate-nickel mixture was then poured into a 20 mL fritted column, and the flowthrough was saved. The remaining resin was then washed with ~20 mL Wash Buffer 1 (50 mM HEPES buffer, pH 7.5), ~20 ml Wash Buffer 2 (50 mM HEPES, 5 mM imidazole, pH 7.5), and eluted with 12 mL Elution Buffer (50 mM HEPES, 300 mM imidazole, pH 7.5). 2-mercaptoethanol (Bio-Rad Laboratories; Hercules, CA, USA) was then added to crude lysate (200 mM final concentration), and the mixture was allowed to incubate with nutation at room temperature overnight. The resulting cleaved protein was then dialyzed against 20 mM Tris pH 8.0 for 8-10 h. The resulting dialysate was then treated with 5 mL nickel agarose resin (GoldBio) and incubated with nutation at 4 °C for 1-2 h. The mixture was then applied to a 20 mL fritted column and flowthrough containing αS was collected in a 15 ml Falcon tube. The resulting enriched protein mixture was then dialyzed against 20 mM Tris, pH 8.0 overnight and purified via FPLC using a 5 ml HiTrap Q-HP column (Cytiva; Marlborough, MA, USA) using the following method: Buffer A: 20 mM Tris, pH 8.0; Buffer B: 20 mM Tris, 1 M NaCl, pH 8.0; Gradient: 0% Buffer B – 5 column volumes, 0-10% Buffer B – 5 column volumes, 20-30% Buffer B – 20 column volumes, 30-100% Buffer B – 10 column volumes; flowrate 3 mL/min. The resulting eluted fractions were then assessed for purity via MALDI-TOF MS, and pure fractions were combined. Protein was then concentrated, and buffer exchanged into phosphate-buffered saline (PBS, NaCl 0.137 M, KCl 0.0027 M, $Na_2HPO_4$ 0.01 M, $KH_2HPO_4$ 0.0018 M) to a final concentration of 100-200 µM via Amicon 3 kDa MWCO filters (Millipore Sigma; St. Louis, MO, USA). Purified protein was aliquoted into 1.5 mL tubes and stored at -80 °C until further use.

**Optimized αS Iridium Labeling.** Each αS cysteine mutant was diluted to 50 µM in PBS in a 15 ml Falcon tube (total vol. 10 ml). TCEP was added (100 µM final conc.), and the resulting solution was allowed to incubate at room temperature for 15-30 minutes. Immediately following incubation, iridium maleimide in DMSO was added from a 20 mM stock (200 µM final conc., 1%DMSO final). The labeling reaction was vortexed briefly and was allowed to rotate at room temperature for 4 hours. Extent of labeling was assessed via MALDI-TOF mass spectrometry. The iridium labeled αS was filtered through a 0.2 µm filter and purified by reverse phase HPLC on a C4 column (gradient 25-60% ACN 0.1% TFA over 32 minutes). The resulting product fractions were combined, diluted 1:1 in PBS to reduce ACN concentration, and dialyzed against PBS overnight. Resulting purified protein was then spin concentrated using Amicon 3.5K MWCO filters prior to aliquoting and freezing until future use.

**Transmission Electron Microscopy (TEM) Imaging.** Samples (5µL) were applied to freshly glow-discharged TEM grids (300-mesh carbon coated copper, Electron Microscopy Sciences) for approx. 1 min and wicked away with filter paper. Excess buffer salts were removed by applying MilliQ water (5µL) to the grids for 30 sec. and then wicking with filter paper. A solution of 2% uranyl acetate in water (5µL) was then applied to grid for 1min and wicked away with filter paper. Samples were then imaged with an accelerating voltage of 120kV and a magnification of 26000x on a Talos 12 TEM (Thermo Fisher).

### Mouse Brain Lysate Preparation and µMap Studies

**Minimal Mouse Brain Lysate Preparation.** Mouse brains were obtained from the Center for Neurodegenerative Disease Research at the University of Pennsylvania. One whole mouse brain was Dounce homogenized in 2 ml of PBS supplemented with EDTA-free Halt protease inhibitors (Thermo Fisher) and divided equally amongst 4 Eppendorf tubes (~500 µl each). The resulting homogenate was then sonicated with a probe sonicator (1 min. amp=50, 2s on 2s off; 1 min. amp=55, 2s on 2s off). Lysate was used directly for subsequent experiments.

**General µMap Irradiation and Western Blotting.** αS (10 µM) and mouse brain lysate (5-10 mg/ml) were combined in an Eppendorf tube (15 µl total vol.) and allowed to incubate at 37°C for 1 hour. Following incubation, 0.15 µl of 25-50mM Dz-BTN (250-500 µM final conc.) was added and samples were transferred to 1 ml clear glass shell vials (Analytical Sales and Services Inc, Flanders, NJ, USA). Samples were then irradiated at 445 nm for 2 minutes with an intensity of 515 mW/well using a Lumidox Controller (Analytical Sales and Services Inc, Flanders, NJ, USA). Following irradiation, glass vials were inverted into an Eppendorf tube and spun briefly at 5000g for 30s to recover biotinylated sample. Cold methanol was added to each sample (4x volume), after which they were incubated at -80C for 1 hour. The samples were then spun down and pelleted at 10000g for 10 minutes, after which the methanol supernatant was removed and allowed to dry completely. The protein pellet was then resuspended in 15µl of PBS and sonicated with a probe sonicator (1 min. amp=50, 2s on 2s off; 1 min. amp=55, 2s on 2s off). 5 µl of 4xLDS+BME (Thermo Fisher, Waltham, MA) was added to each sample, and the samples were heated at 75°C for 10 minutes. Samples were then subjected to SDS-PAGE using a 1.0 mm 4-12% NuPAGE Bis-Tris gel (Thermo Fisher, Waltham, MA), and were immediately transferred to a 0.20 µm Nitrocellulose membrane via wet-transfer western blot. Efficient transfer was analyzed via visualization with Ponceau red stain and the blot was visualized using either fluorescent AlexFluor conjugated antibodies or streptavidin-HRP fusion and enhanced chemiluminescent substrate. (Thermo Fisher, Waltham, MA)

**µMap Fluorescence SDS-PAGE.** αS (10 µM) and mouse brain lysate (5-10 mg/ml) were combined in an Eppendorf tube (15 µl total vol.) and allowed to incubate at 37°C for 1 hour. Following incubation, 0.15 µl of 5-10mM **Dz-TMR** (50-100 µM final conc.) was added and samples were transferred to 1 ml clear glass shell vials (Analytical Sales and Services Inc, Flanders, NJ, USA). Samples were then irradiated at 445 nm for 2 minutes with an intensity of 515 mW/well using a Lumidox Controller (Analytical Sales and Services Inc, Flanders, NJ, USA). Following irradiation, glass vials were inverted into an Eppendorf tube and spun briefly at 5000g for 30s to recover biotinylated sample. 5 µl of 4xLDS+BME (Thermo Fisher, Waltham, MA) was added to each sample, and the samples were heated at 75°C for 10 minutes. Samples were then subjected to SDS-PAGE using a 1.0 mm 4-12% NuPAGE Bis-Tris gel (Thermo Fisher, Waltham, MA), and were immediately analyzed via multiplexed fluorescence imaging using a G:BOX mini gel imager (Syngene, Cambridge, UK).

**µMap Interactome Protein Isolation.** αS (10 µM) and mouse brain lysate (20 mg/ml) were combined in an Eppendorf tube (50 µl total vol.) and allowed to incubate at 37°C for 1 hour. Following incubation, 0.50 µl of 50mM **Dz-DTB** or **Dz-BTN** (500 µM final conc.) was added and the samples were transferred to 1 ml clear glass shell vials (Analytical Sales and Services Inc, Flanders, NJ, USA). Samples were then irradiated at 445 nm for 2 minutes with an intensity of 515 mW/well using a Lumidox Controller (Analytical Sales and Services Inc, Flanders, NJ, USA). Following irradiation, glass vials were inverted into an Eppendorf tube and spun briefly at 5000g for 30s to recover desthiobiotinylated or biotinylated sample. 950 µl of PBS was added to each tube, and the samples were centrifuged (16,100 g) for 1 hr. The soluble protein fraction was collected, added directly to streptavidin agarose resin (50 µl bed volume) (GoldBio; St. Louis, MO, USA), and incubated overnight at 4°C. After overnight incubation, the beads were washed 4 times with PBS and once with $H_2O$ by centrifuging (0.5 min, 0.5 g) and removing the supernatant. The beads were then incubated with 500 µl of biotin (2 mM final conc.) in $H_2O$ for 1-2 hours. After incubation, the beads were pelleted (0.5 min. 0.5 g) and the supernatant was collected. The beads were then washed with a second aliquot of 500 µl biotin (2 mM final conc.), and the supernatants were combined, and spin concentrated to 100 µl using Amicon 3.5 kD MWCO filters (Thermo Fisher, Waltham, MA). The resulting eluted proteins were then flash frozen in liquid nitrogen and concentrated using a speed vacuum, resulting in a lyophilized protein pellet.

**MS Sample Digest and Preparation.** A lyophilized protein pellet from µMap interactome studies was denatured and reduced by addition of 20 µl of urea (8M final conc.) and 0.8 µl dithiothreitol (DTT) (20 mM final conc. from 500 mM stock prepared fresh) with heating at 50 °C. The sample was allowed to cool to room temperature, and free cysteines were alkylated by addition of 2 µl iodoacetamide (IAA) (40 mM final conc. from 400 mM stock prepared fresh) for 30 min. in the dark. The remaining IAA was quenched by addition of another 0.8 µl of DTT (40 mM final conc., e.g., 20 mM on top of the 20 mM previously added). To each reaction 20 µl of $NH_4HCO_3$ (500 mM, pH = 8.0) was added followed by 160 µl of $H_2O$ ($NH_4HCO_3$ final conc. = 50 mM). 1.0 µg of trypsin was added to each sample, and digestions were allowed to proceed at 37°C overnight. After digestion samples were acidified by addition of 2.0 µl trifluoroacetic acid (TFA) (1% final conc.), and the samples were cleaned up using Pierce C18 spin columns (Thermo Fisher, Waltham, MA). The resulting peptide mixtures were concentrated using a speed vacuum and were stored as a lyophilized pellet until analysis by MS.

**MS Analysis.** Digested samples were analyzed by a Q-Exactive HF mass spectrometer (Thermo Fisher Scientific) coupled to a Dionex Ultimate 3000 UHPLC system (Thermo Fischer Scientific) equipped with an in-house made 15 cm long fused silica capillary column (75 µm ID), packed with reversed phase Repro-Sil Pure C18-AQ 2.4 µm resin (Dr. Maisch GmbH, Ammerbuch, Germany). Elution was performed by the following method: a linear gradient from 4 to 38% buffer B (90 min), followed by 95% buffer B (5 min), and re-equilibration from 95 to 4% buffer B (5 min) with a flow rate of 300 nL/min (buffer A: 0.1% formic acid in water; buffer B: 80% acetonitrile with 0.1% formic acid). Data were acquired in data-dependent MS/MS mode. Full scan MS settings were as follows: mass range 200–1600 m/z, resolution 120,000; MS1 AGC target 3E6; MS1 Maximum IT 100. MS/MS settings were: resolution 30,000; AGC target 5E5; MS2 Maximum IT 100 ms; fragmentation was enforced by higher-energy collisional dissociation with stepped collision energy of 25, 27, 30; loop count top 20; isolation window 1.4; MS2 Minimum AGC target 800; charge exclusion: unassigned, 1, 8 and >8; peptide match preferred; exclude isotope on; dynamic exclusion 45 s.

**Data Analysis for Radii Determination.** Samples were searched using MaxQuant (Version 2.1.4.0) with a standard label-free quantitation workflow and **Dz-DTB** included as a modification on any amino acid. A library of identified covalently labeled peptides was assembled for $\alpha$S-$C^{Ir}_{12}$, $\alpha$S-$C^{Ir}_{62}$, and $\alpha$S-$C^{Ir}_{114}$ and a monomeric ensemble (n=1000) of αS was generated using Fast Floppy

Tail in PyRosetta.[4] A theoretical distance ($C_\alpha$- $C_\alpha$) was calculated from the position of iridium labeling to each experimentally identified modified residue, for each member of the ensemble. The average radius was then computed from the ensemble distance list to approximate the radius of labeling based on experimentally obtained photolabeling data.

**Data Analysis for αS-Ir Interactomes.** Samples were searched in Spectronaut against mouse reference sequence database from UniProt. Analysis was conducted using RStudio 4.3.2 for Windows. Values that are missing not at random (MNAR) were accounted for by removing protein values that were not identified in all replicates of at least one condition. After filtering, individual samples were normalized by median centering. Missing values were imputed using the 'manual_impute' function of the Differential Enrichment analysis of Proteomics data (DEP) downloaded from CRAN 4.5.0. Differential abundance analysis was used using limma 3.64.3 package from Bioconductor 3.21 to generate pair-wise comparisons between conditions, with p-values adjusted using Benjamini-Hochberg correction. Over representation analysis (ORA) was done with clusterProfiler 4.16.0 from Bioconductor 3.21 using p-value adjusted protein hits with the nonfiltered protein list as the background set. Our µMap interactome dataset in mouse brain lysate was also compared to APEX-based PL systems in human Lewy Bodies and rat cortical neurons. Open-source proteomic files were downloaded from each study ("NIHMS872631-supplement-Supp_Table_1", "pnas.2114405119.sd01"). An API to the NCBI database was created using retrenz 1.2.4 package, after which all non-mouse proteins were converted into FASTA files containing their amino acids sequences. The whole mouse proteome was downloaded open-source from UniProt to create a local database for BLAST alignment to get homologues for subsequent analysis. Shared hits are defined as the non-mouse protein homologues and native mouse proteins found in both µMap and APEX PL studies.

**Primary Cell Culture and αS Treatment.** All procedures were approved by the University of Pennsylvania Institutional Animal Care and Use Committee and in accordance with the experimental guidelines outlined by the National Institutes of Health Guide for the Care and Use of Experimental Animals. Primary neonate neurons were harvested from the hippocampus of embryonic day E16-E18 CD1 mouse embryos. The dissociated neurons were plated on a Perkin Elmer 96W ViewPlate at a density of 17,500 cells/well in neurobasal media (Thermo Fisher) supplemented with B27 (Thermo Fisher), 2mM GlutaMax (Thermo Fisher), and 100 U/mL penicillin/streptomycin (Thermo Fisher) with 5% Fetal Bovine Serum (Thermo Fisher). On DIV 1, the media was aspirated off and replaced with 100µL of FBS-free media. On DIV 7, recombinant αS Ir and WT αS stocks were sonicated (Diogenode Bioruptor UCD-300 bath sonicator) for 20 cycles (30s on, 30s off), after which the plate was treated with αS for a final concentration of 47 ng/µL and 13 ng/µL of monomers and fibrils respectively per condition. The treated plates were left to incubate overnight at 37 °C.

**Hippocampal µMap Interactome by Immunochemistry.** Following overnight incubation, 250µM **Dz-BTN** affinity label was added to each well from a 100mM stock in DMSO and placed back in the incubator for 15 minutes. Wells that were not probed with Dz-BTN were similarly treated with a DMSO vehicle. The cells were then washed with neurobasal media to remove extracellular αS and Dz-BTN and exchanged into PBS prior to irradiation. The plates were irradiated at 445 nm at 515mW/well for 2 minutes using a Lumidox Controller (Analytical Sales and Services Inc, Flanders, NJ, USA). The plates were fixed with a solution of 4% paraformaldehyde and 4% sucrose and washed with PBS, and the cells were permeabilized with

1% Triton-X100 and 0.5% Tween 20 in PBS. Blocking Buffer for Fluorescent Western Blotting (Rockland Immunochemicals) was added, and the plates were incubated at room temperature for 1 hr, followed by incubation with primary antibodies overnight at 4°C (Table S1). After incubation, the wells were washed 3 times with PBS and incubated with fluorescent secondary antibodies and Alexa Fluor 555 streptavidin for 1 hr at room temperature. The wells were washed again with PBS and stained with a DAPI solution (Thermo Fisher, 1:10,000 dilution in DPBS). The plates were sealed with adhesive covers and scanned with an In Cell Analyzer 2200 (GE Healthcare) with a 40x objective. The respective channels were imaged and analyzed with Image J.

**DNA Paint Immunostaining.** Dissociated hippocampal neurons were plated on an ibiTreat µ-Slide 8 Well chamber slide (Ibidi) at a density of 20,000 cells/well in neurobasal media. Standard cell culture protocol as described above was followed until DIV 7, after which the wells were treated with either 0, 50, or 100 ng of monomeric WT αS. Cells were fixed with a solution of 4% paraformaldehyde and 4% sucrose, washed with PBS, and permeabilized with 1% Triton-X100 and 0.5% Tween 20 in PBS. The cells were then blocked for 1 hour at room temperature while shaking with primary blocking buffer (10% donkey serum, 0.1% saponin, 0.05 mg/mL salmon sperm DNA) in PBS. Cells were then incubated overnight at 4°C with primary antibodies diluted in blocking buffer as indicated in **Table S1**. Following primary antibody incubation, cells were washed with PBS 3x for 5 min each, and Massive Photonics washing buffer 1x for 5 minutes at room temperature. Cells were then incubated with the secondary antibodies diluted in antibody incubation buffer (Massive Photonics) as indicated in SI XX for 2 hours at room temperature and shaking. Finally, cells were washed with PBS 3x for 5 min each, and Massive Photonics washing buffer 1x for 5 minutes at room temperature and stored at 4°C until imaging.

**DNA-PAINT Imaging.** DNA-PAINT imaging was captured using the ONI Nanoimager at 30°C using HiLo illumination. Laser power was set at 15 mW for the 561 and 640 laser channels. Appropriate imager oligomer strands solutions were developed at a concentration of 0.5nM in imaging buffer (Massive Photonics) and added to the imaging chambers. One imager is conjugated with Cy3B and the other imager is conjugated with ATTO655 for dual-color imaging. The samples were imaged at 100-ms exposure using a laser program that captures a total of 50,000 frames, and 25,000 frames per target. Images were exported from NimOS software with the filtering parameters being a localization precision of 30 nm, and a Sigma X/Y between 10 and 250 nm. Analysis using MATLAB R2025a custom made code[5] determined 18.2% of mitochondrial structures labeled with GLUD1 are enriched with αS localizations.

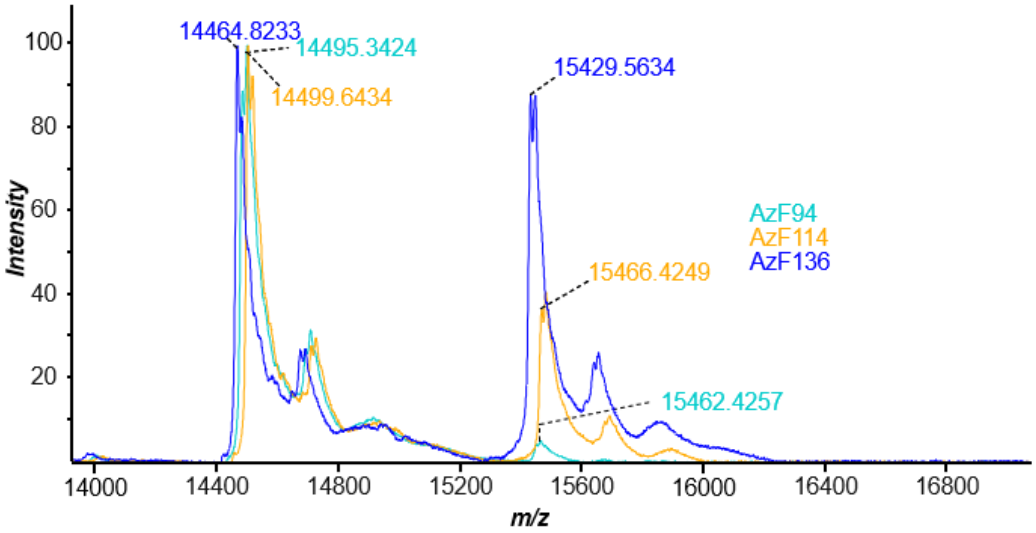


**Figure S1.** The efficiency of Iridium attachment by copper-free αS-AzF constructs and TAMRA-DBCO as visualized by MALDI-TOF mass spectrometry. Masses of 14465.8-14499.6434 m/z denotes unclicked αS-AzF constructs for F94, F114, F136 mutants. Second local maxima with masses 15429.5634-1562.4257 m/z shows successful TAMRA-DBCO conjugation, with ~50% relative conversion.

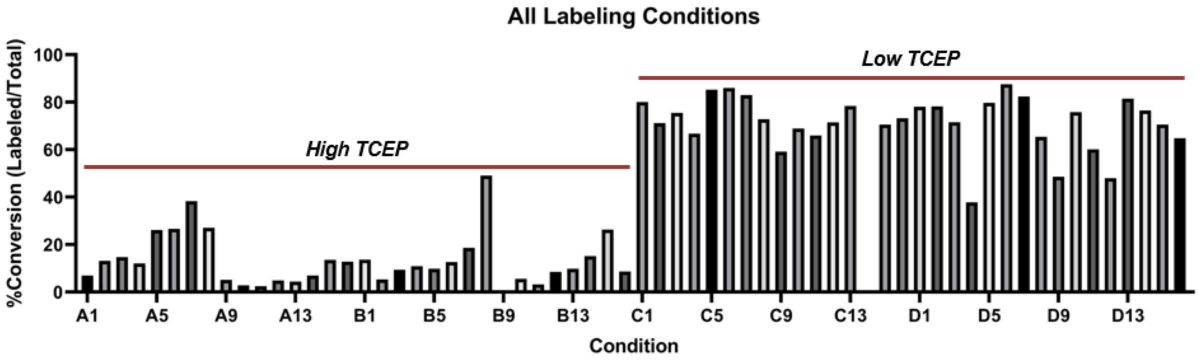


**Figure S2.** Optimized protein and TCEP concentrations for Ir-maleimide labeling

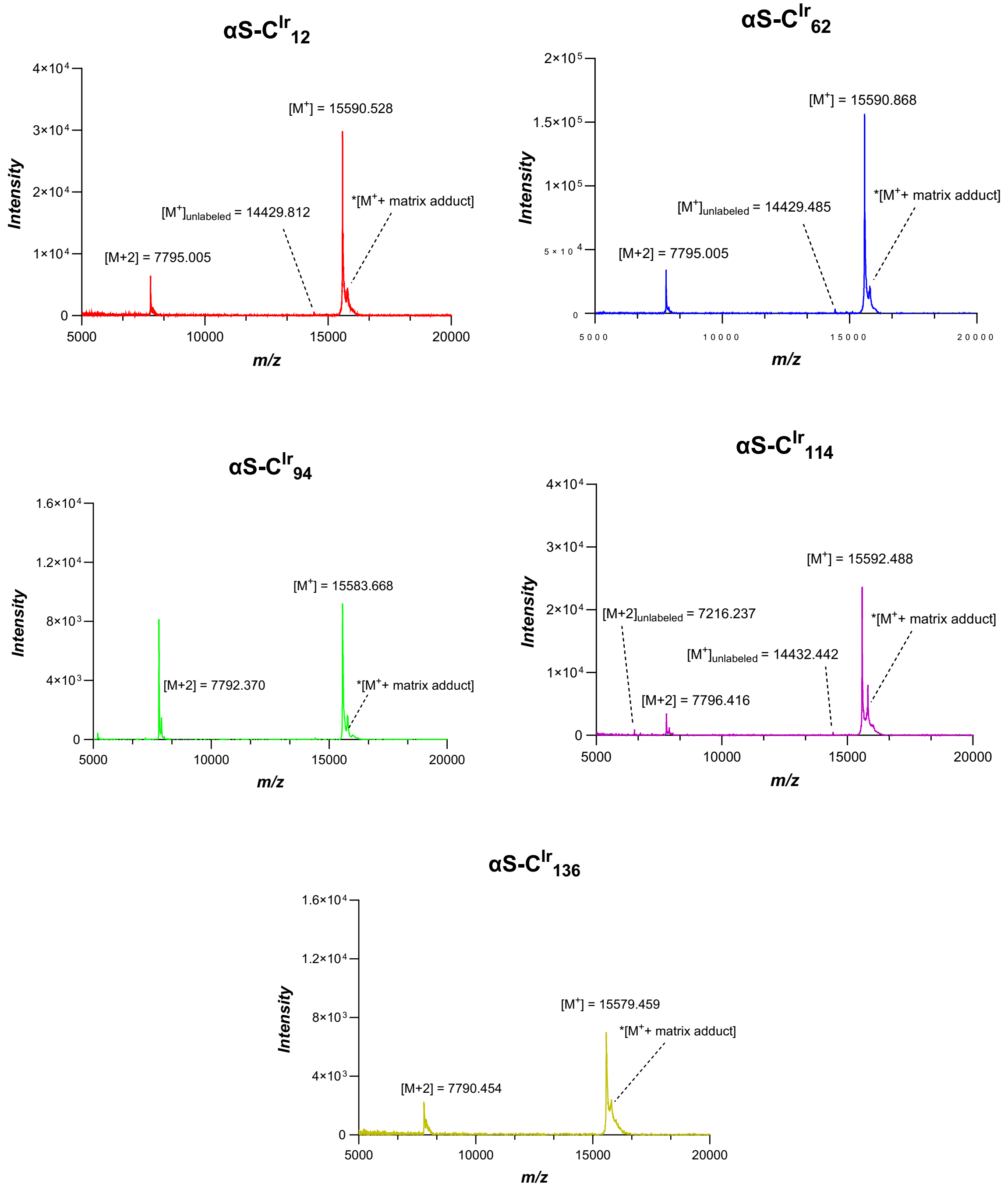


**Figure S3.** MALDI-TOF mass spectrometry after HPLC purification shows pure fractions of αS-$C^{Ir}{}_{12}$, αS-$C^{Ir}{}_{62}$, αS-$C^{Ir}{}_{94}$, αS-$C^{Ir}{}_{114}$, αS-$C^{Ir}{}_{136}$ constructs. Molecular $[M^+]$ ion peak denotes Iridium-labeled αS constructs, along with [M+2] peak. *$[M^+$ + matrix adduct] peak denotes labeled construct with Sinapic acid matrix (+224.21 m/z). Trace amounts of unlabeled αS-$C_n$ constructs are indicated as $[M^+]_{unlabeled}$.

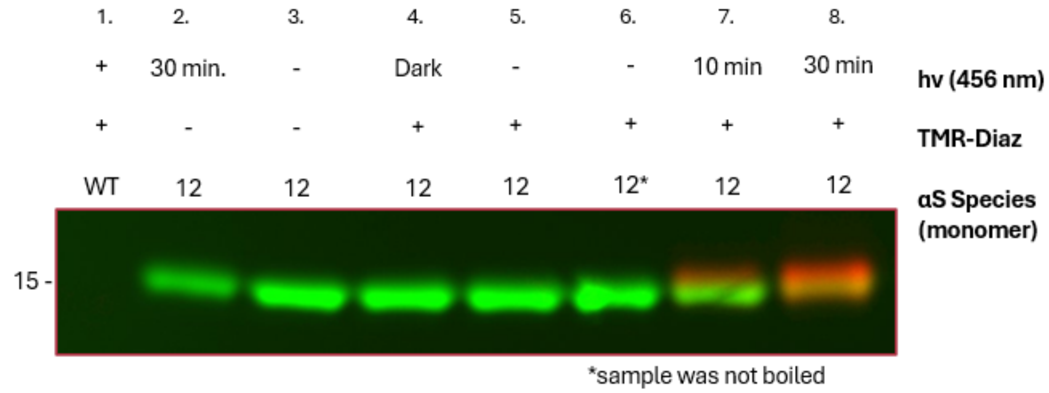


**Figure S4.** Protein photo-conjugation is dependent on the presence of a diazirine species, Iridium catalyst, and blue light. αSIr was irradiated in the presence of Dz-TAMRA, after which self-crosslinking was determined by gel for both Iridium fluorescence from monomeric αS-$C^{Ir}_{12}$ (green) and TAMRA fluorescence (red). Both signals absent with WT αS (Lane 1). Samples were irradiated using a 456nm PR160 Kessil Lamp for either 10 minutes (Lane 7) or 30 minutes (Lane 2, Lane 8). All samples were worked up under normal laboratory lighting except for Lane 4, which was worked up under red light to test background diazirine reactivity under ambient light. To test the effect of SDS boiling on Iridium fluorescence, sample in Lane 6 was not boiled and loaded directly under the gel.

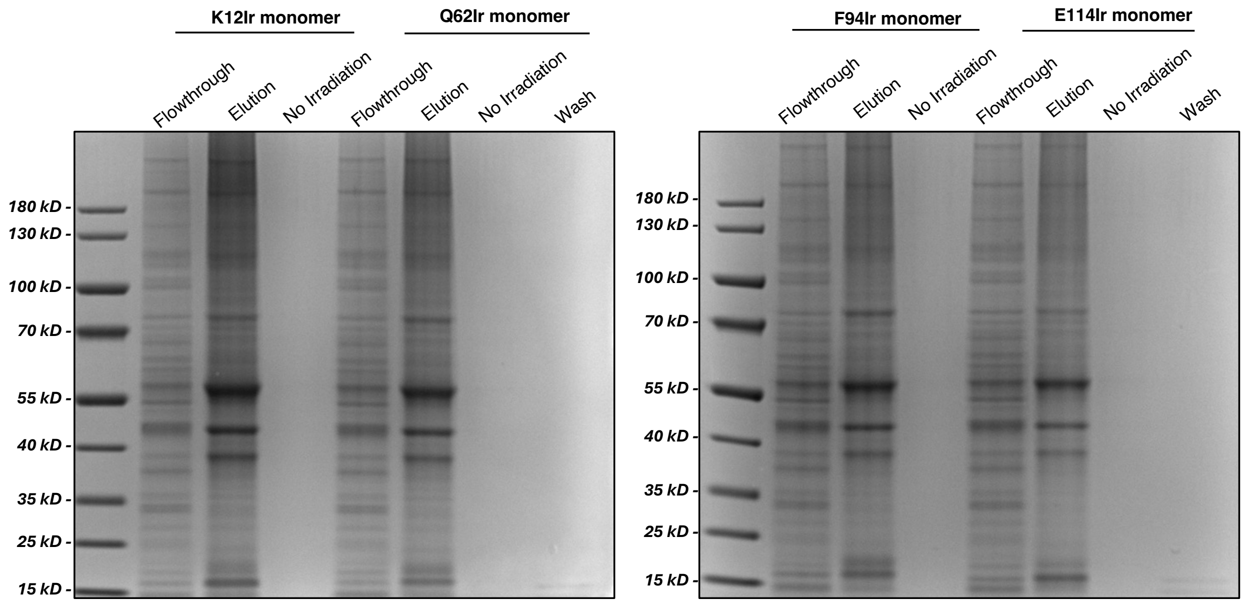


**Figure S5.** Gels showing enrichment of specific proteins for αS-C$^{Ir}$ constructs using Dz-DTB followed by biotin elution.

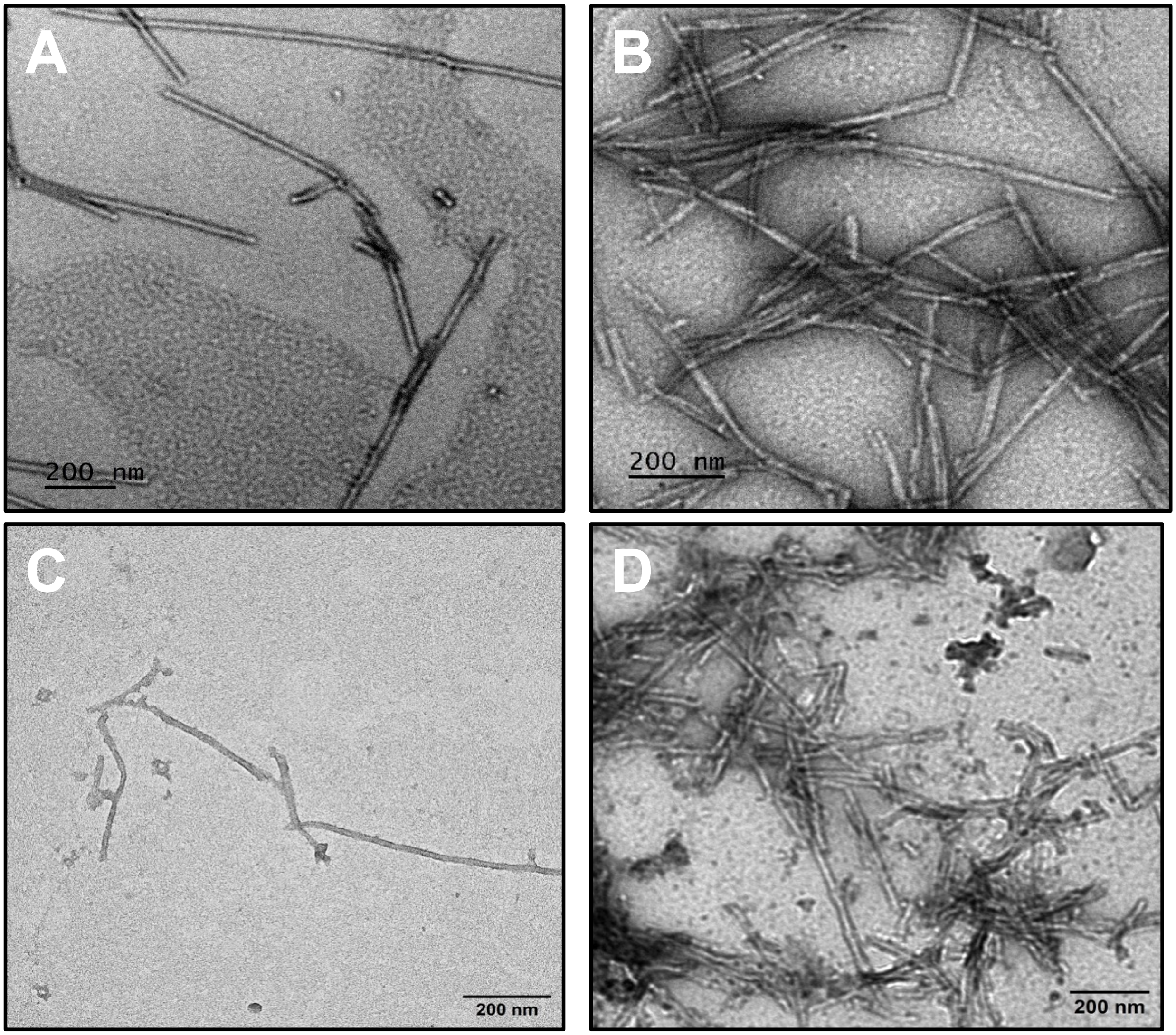


**Figure S6.** TEM images of (A) WT αS PFFs, (B) 10% αS-$C^{Ir}_{12}$ + 90% WT αS PFF mix, (C) 100% αS-$C^{Ir}_{62}$ PFFs, and (D) 100% αS-$C^{Ir}_{114}$ PFFs. Monomeric constructs were aggregated at 100μM for 72 hours at 37°C while shaking at 350rpm. Samples were not centrifuged to remove monomeric fraction before imaging to avoid significant fibril clumping. αS-$C^{Ir}_{62}$ PFFs showed notably less PFF formation at the 3-day mark in comparison to other constructs, suggesting NAC region attachment perturbs aggregation kinetics.

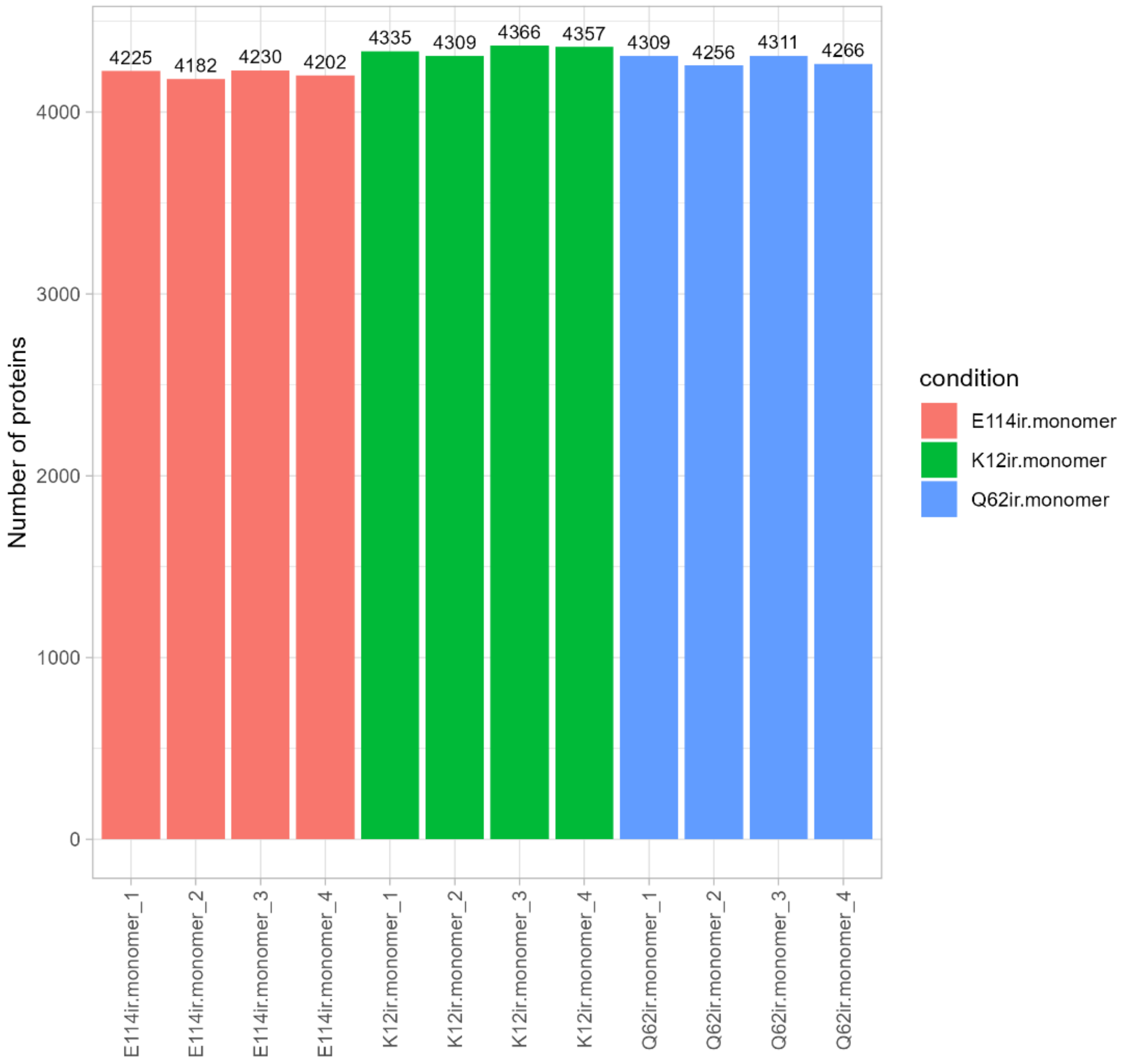


**Figure S7.** Bar plot of biotin-conjugated proteins labeled in mouse brain lysate doped with monomeric αS-$C^{Ir}_{12}$, αS-$C^{Ir}_{62}$, and αS-$C^{Ir}_{114}$. Following blue light irradiation in the presence of Dz-BTN, enWriched lysate was incubated with Streptavidin beads that were then subjected to tryptic digest and followed by MS analysis of photocrosslinked proteins. Plot represents proteins found in all replicates of at least one labeled monomeric position (ie. αS-$C^{Ir}_{12}$), filtering out MNAR values below the detection limit.

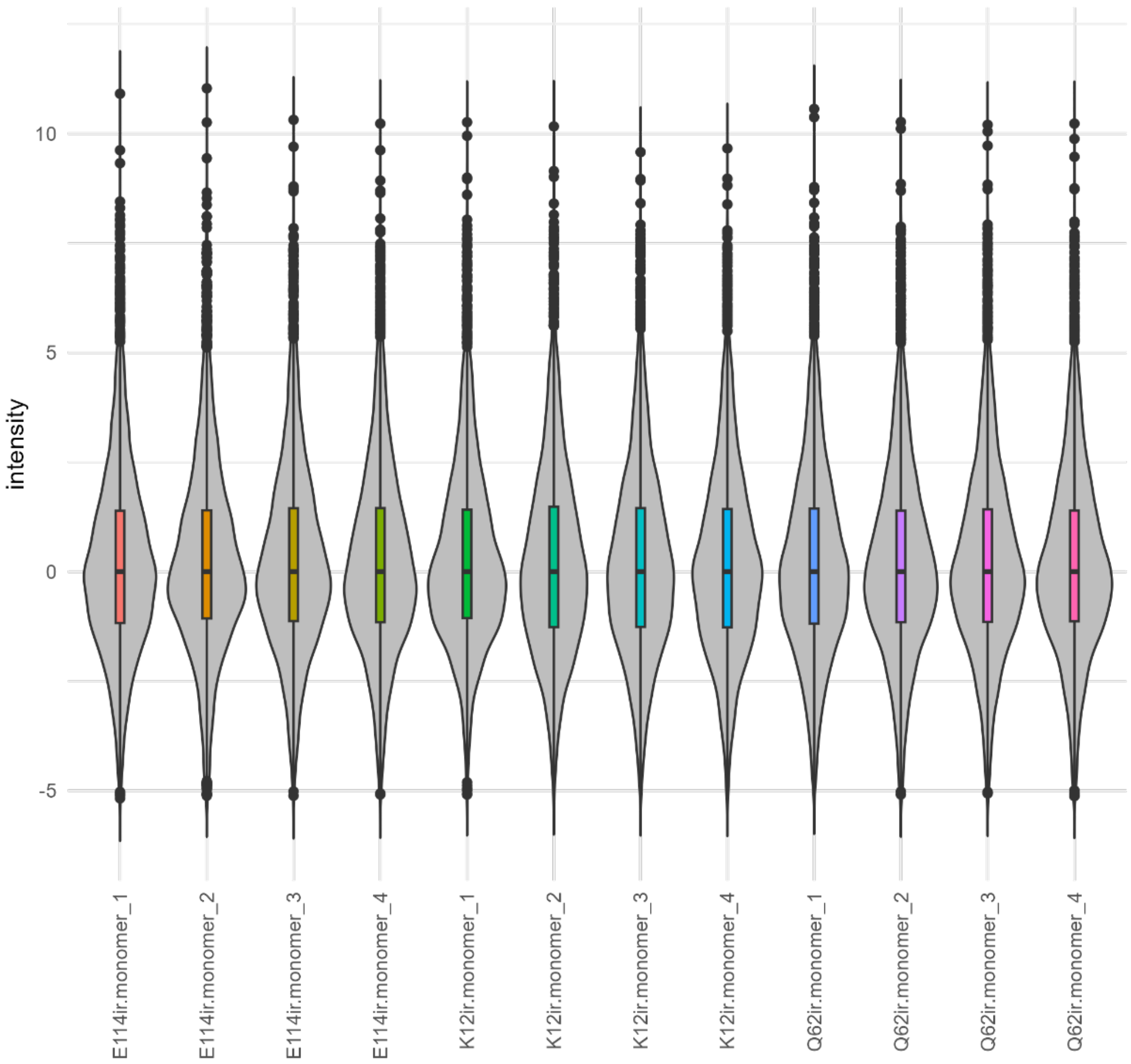


**Figure S8.** Violin and boxplot of normalized protein intensities after data filtering in αS-$C^{Ir}_{12}$, αS-$C^{Ir}_{62}$, and αS-$C^{Ir}_{114}$ monomer-doped lysate samples. Intensities were normalized and scaled to center individual sample medians (median = 0).

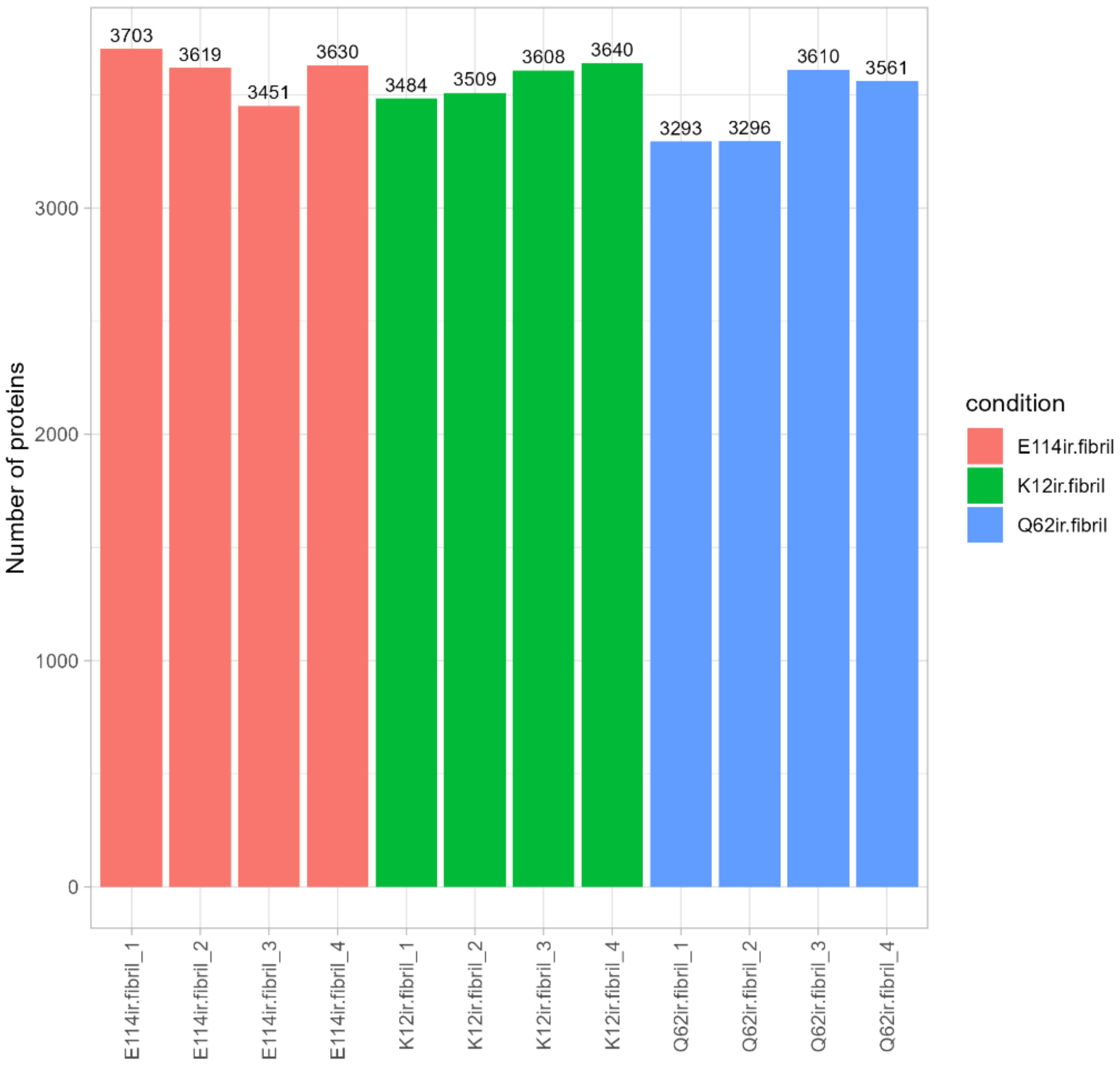


**Figure S9**. Bar plot of biotin-conjugated proteins labeled in mouse brain lysate doped with αS-$C^{Ir}{}_{12}$, αS-$C^{Ir}{}_{62}$, and αS-$C^{Ir}{}_{114}$ PFF. Plot represents proteins found in all replicates of at least one labeled PFF position, filtering out MNAR values below the MS detection limit after Dz-BTN photoconjugation of interactome proteins in spiked mouse brain lysate.

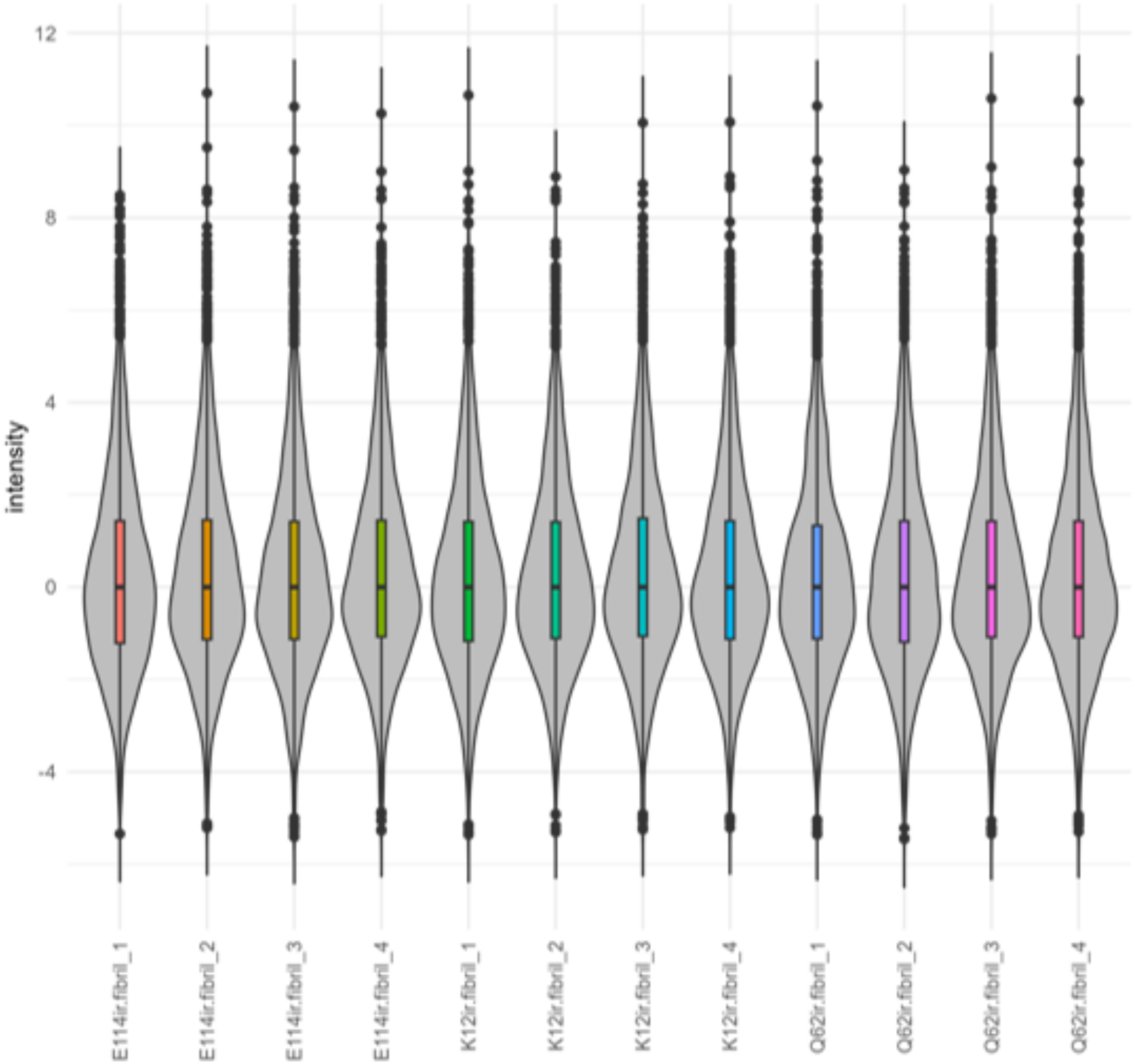


**Figure S10.** Violin and boxplot of normalized protein intensities after data filtering in αS-$C^{Ir}_{12}$, αS-$C^{Ir}_{62}$, and αS-$C^{Ir}_{114}$ PFF-doped lysate samples. Intensities were normalized and scaled to center individual sample medians (median = 0).

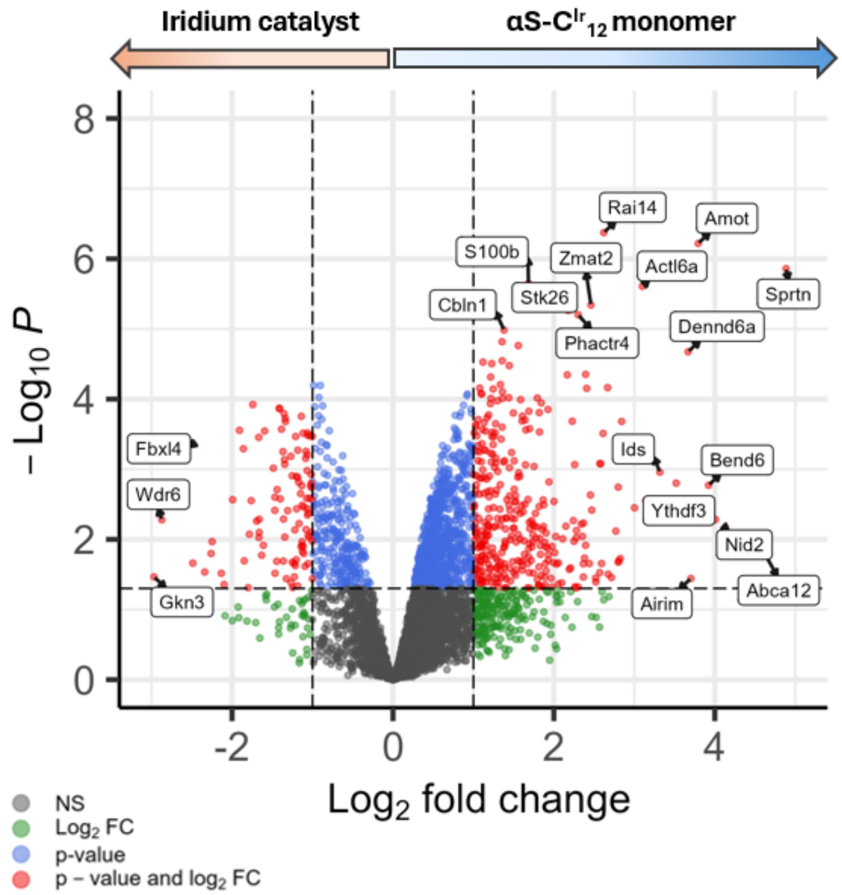


**Figure S11.** Volcano plot of biotin-conjugated mouse brain proteins enriched by free Iridium catalyst (negative $Log_2$ fold-change values) versus αS-$C^{Ir}_{12}$ monomer (positive $Log_2$ fold-change values). Free Iridium catalyst used as an over-enrichment control and a measure of promiscuous labeling. y = 1 denotes threshold where there is two-fold enrichment of proteins in αS-$C^{Ir}_{12}$ monomer condition over Iridium catalyst only. Horizontal axis at x = 1.313 denotes threshold where -$Log_{10}$(P) meets the threshold adjusted p-value = 0.05.

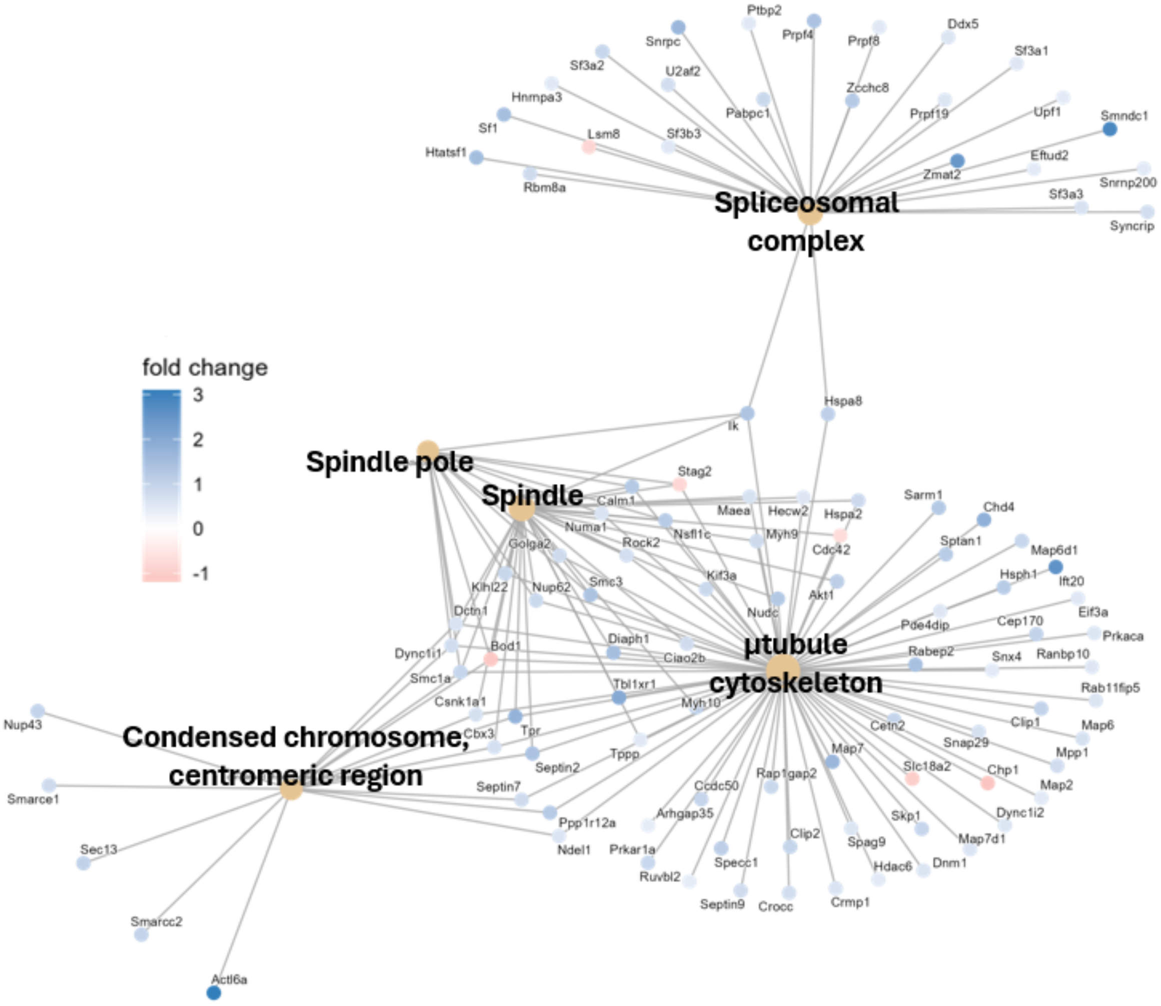


**Figure S12.** Over-representation analysis (ORA) of bioCtin-conjugated mouse brain proteins enriched by either αS-$C^{Ir}_{12}$ monomer or free Iridium catalyst above the adjusted p-value cutoff of 0.05. Gene Ontology (GO) used to cluster proteins into cellular components and biological processes. Blue protein nodes indicate over-enrichment by αS-$C^{Ir}_{12}$ monomer on a $\log_2$ fold change scale while red nodes indicate over-enrichment by free catalyst.

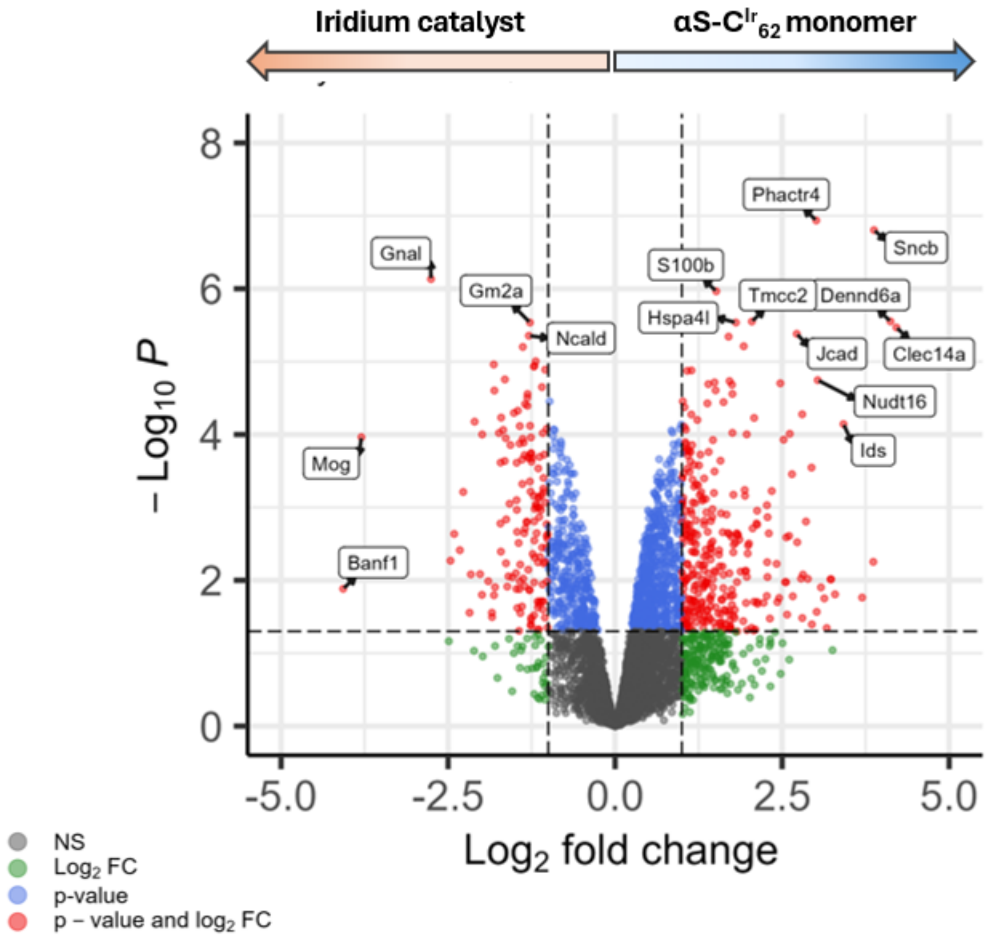


**Figure S13.** Volcano plot of biotin-conjugated mouse brain proteins enriched by free Iridium catalyst (negative $Log_2$ fold-change values) versus αS-$C^{Ir}_{62}$ monomer (positive $Log_2$ fold-change values). Free Iridium catalyst used as an over-enrichment control and a measure of promiscuous labeling. y = 1 denotes threshold where there is two-fold enrichment of proteins in αS-$C^{Ir}_{62}$ monomer condition over Iridium catalyst only. y = 1 denotes threshold where there is a two-fold enrichment from Iridium catalyst. Horizontal axis at x = 1.313 denotes threshold where -$Log_{10}$(P) meets the threshold adjusted p-value = 0.05.

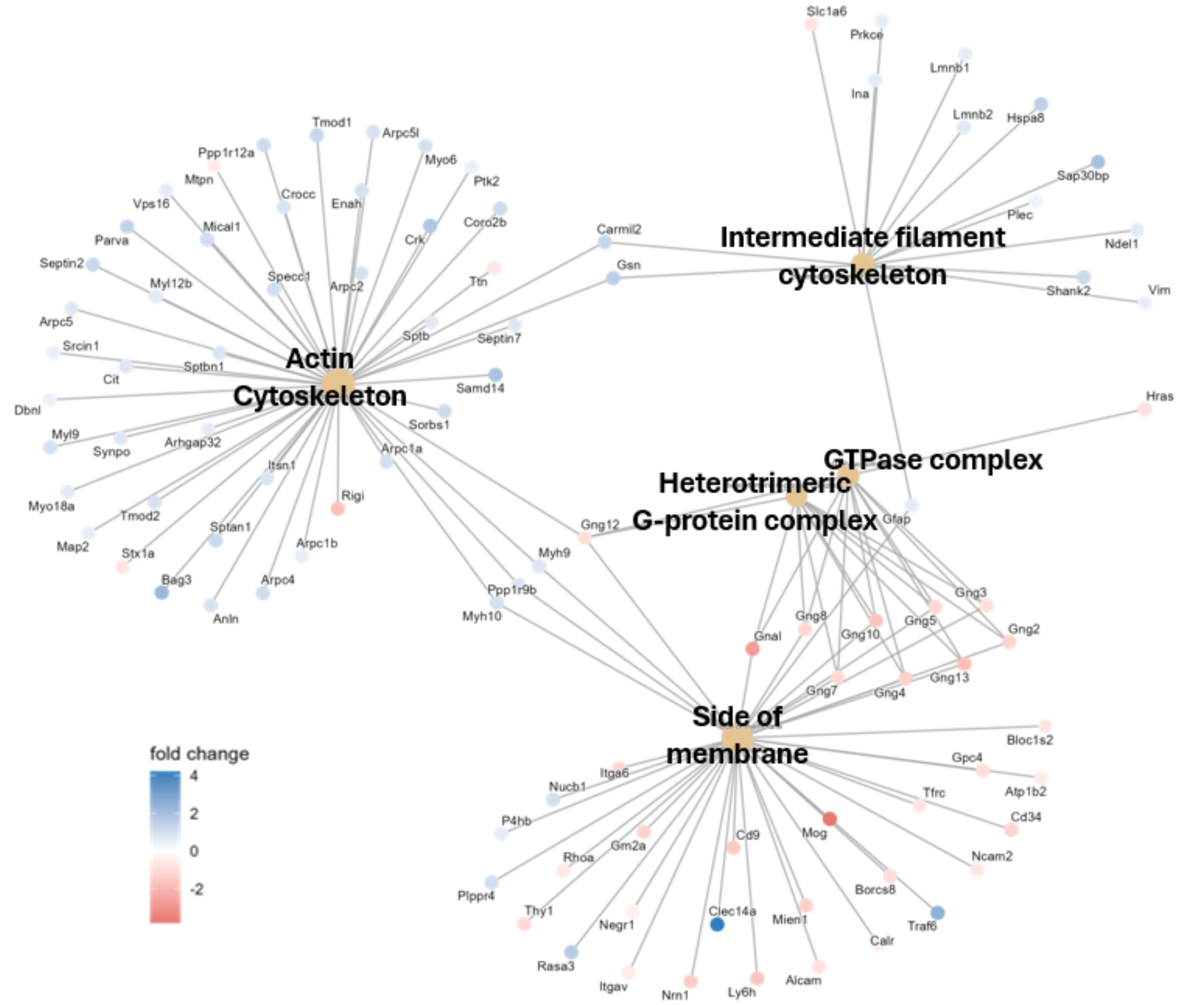


**Figure S14.** ORA of enriched mouse brain proteins by either αS-$C^{Ir}_{62}$ monomer or free Iridium catalyst. Statistically significant hits above p-value = 0.05 were used for GO analysis to cluster proteins into cellular components. Blue protein nodes indicate over-enrichment by αS-$C^{Ir}_{62}$ monomer on a $\log_2$ fold change scale while red nodes indicate over-enrichment by free catalyst, which is used as a measure of promiscuous labeling.

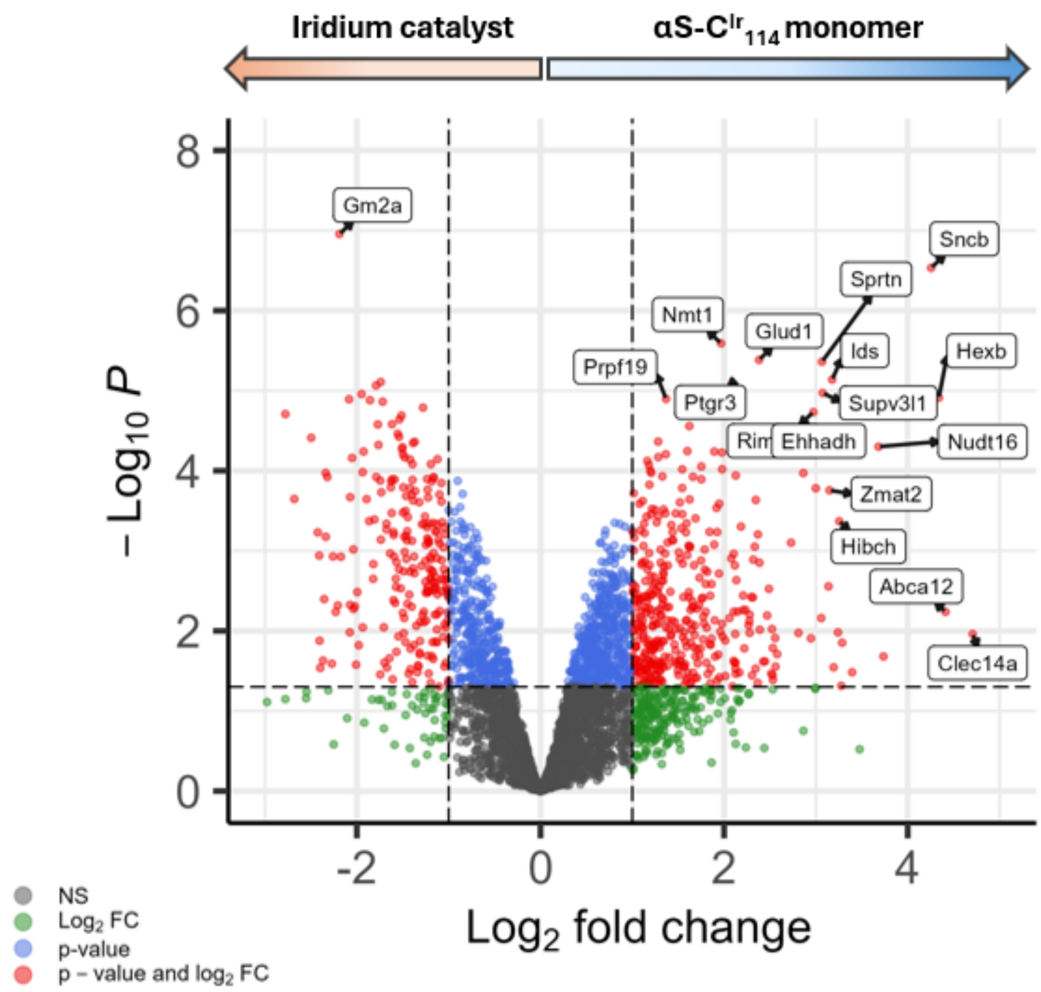


**Figure S15.** Volcano plot of biotin-conjugated mouse brain proteins enriched by free Iridium catalyst (negative $Log_2$ fold-change values) versus αS-$C^{Ir}_{114}$ monomer (positive $Log_2$ fold-change values). Free Iridium catalyst used as an over-enrichment control and a measure of promiscuous labeling.

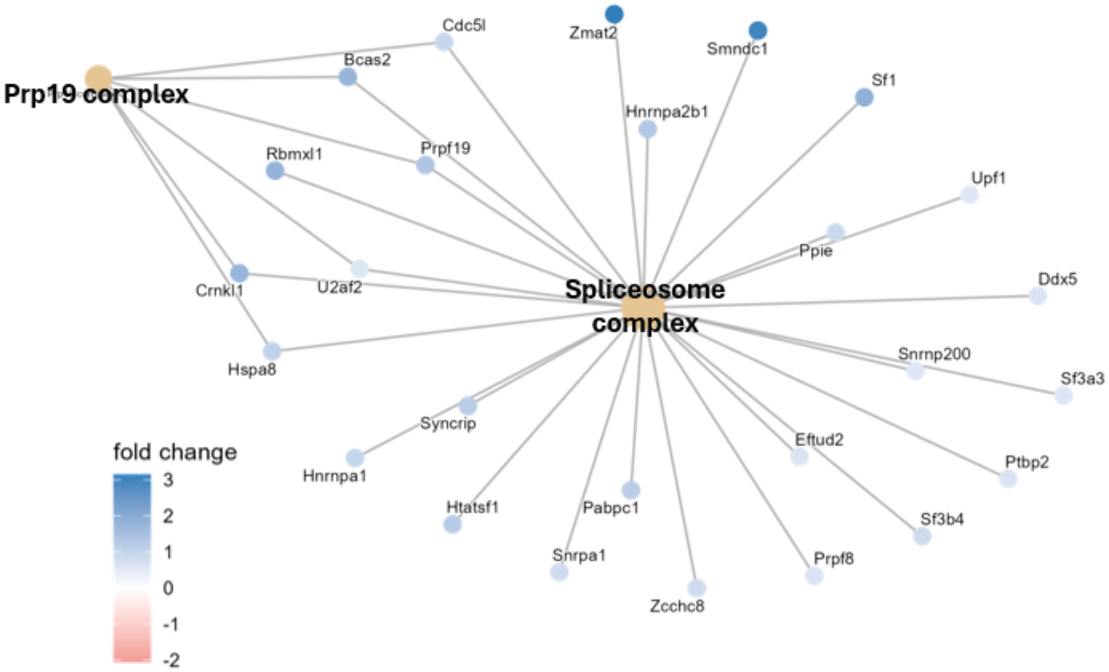


**Figure S16.** GO network-node chart for statistically significant (adjusted p-value = 0.05) mouse brain lysate proteins enriched by αS-$C^{Ir}_{114}$ monomer versus that by free catalyst. Scale bar color represents $\log_2$ fold change in intensity, where blue nodes represent over-representation in αS-$C^{Ir}_{114}$ monomer condition.

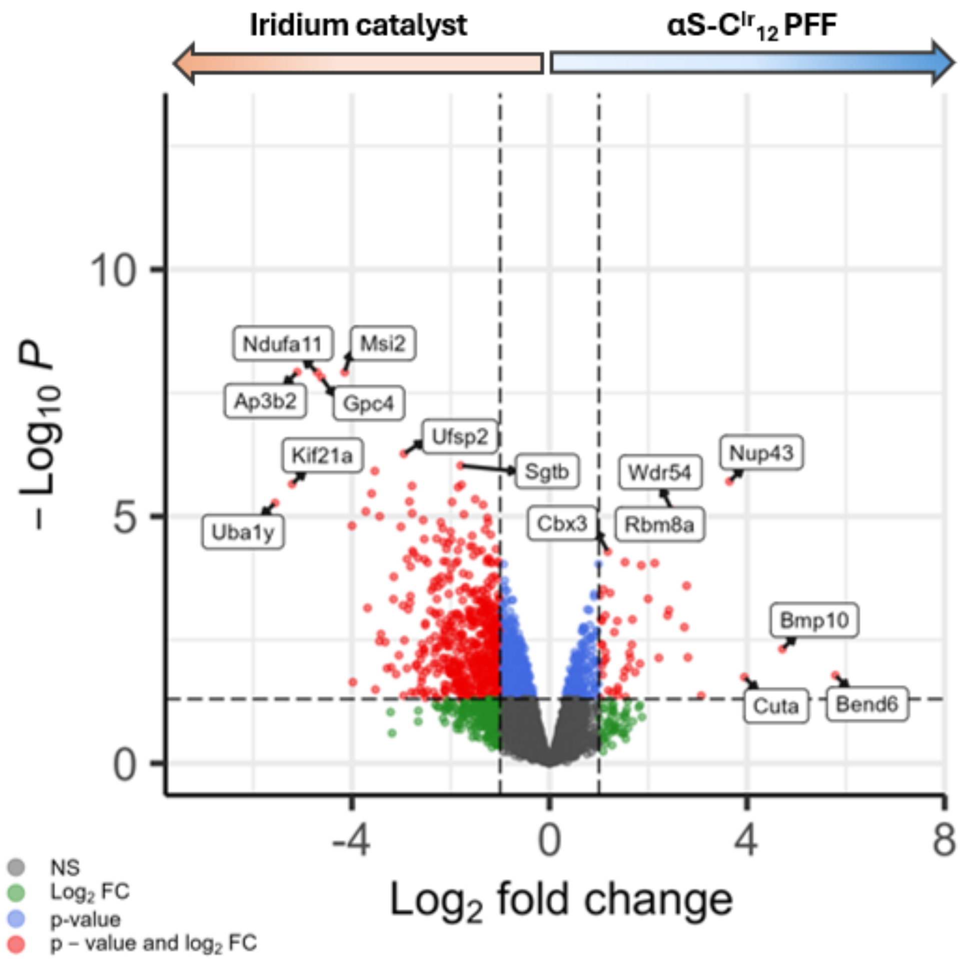


**Figure S16.** Biotin-conjugated mouse brain lysates with negative $\text{Log}_2$ fold change values indicate enrichment by free Iridium catalyst versus positive values for enrichment by αS-$C^{Ir}_{12}$ PFF. Horizontal axis -$\text{Log}_{10}$P = 1.313 indicates threshold of adjusted p-value = 0.05. Labeled proteins on the top right quadrant indicate enriched lysate proteins that are both statistically significant and with a >2 fold change intensity from the promiscuous labeling of the Iridium catalyst condition.

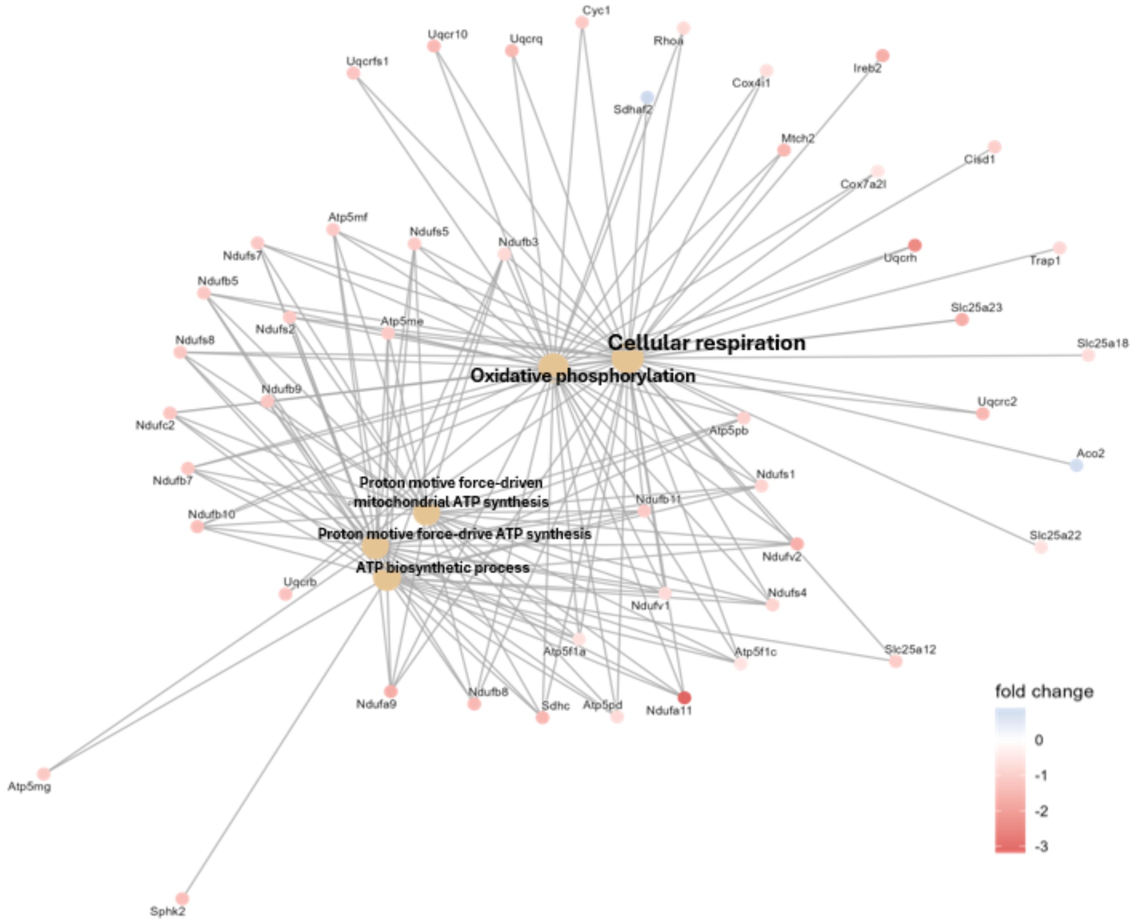


**Figure S17.** GO plot of statistically significant protein hits (adjusted p-value < 0.05) from αS-$C^{Ir}_{12}$ PFF versus Iridium catalyst. Red nodes indicate lysate proteins that are more enriched by Iridium catalyst according to scale color. Clustering indicates components biased toward essential cellular processes like respiration and ATP-synthesis due to overrepresentation of abundant proteins enriched by promiscuous Iridium catalyst labeling.

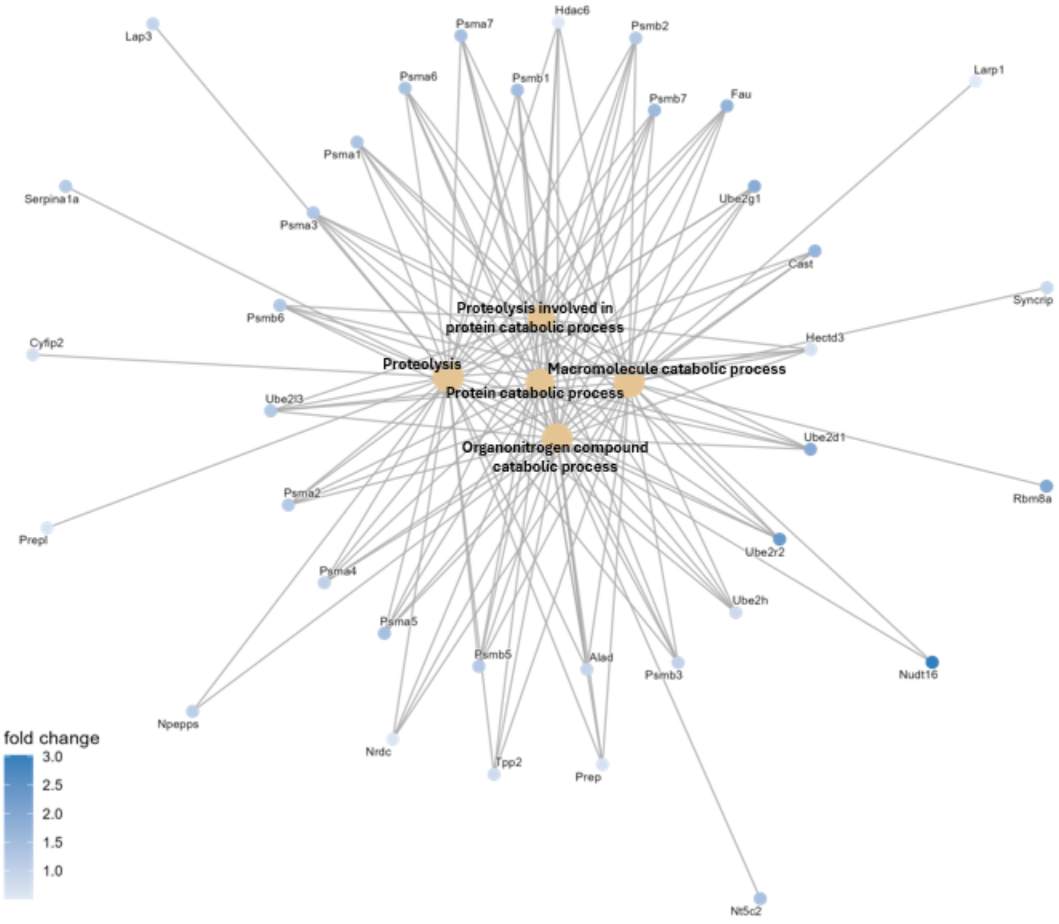


**Figure S18.** GO plot of statistically significant protein hits only from αS-$C^{Ir}_{12}$ PFF enrichment. Out of all statistically significant protein hits (adjusted p-value $< 0.05$) from pair-wise comparison of αS-$C^{Ir}_{12}$ PFF enriched lysate proteins versus Iridium catalyst enriched proteins, only the positive fold change terms (proteins enriched by αS-$C^{Ir}_{12}$ PFF) were taken for ORA clustering. Only blue nodes representing these αS-$C^{Ir}_{12}$ PFF enriched terms are visualized in the resultant network-node graph on a color scale based on $\log_2$ fold change.

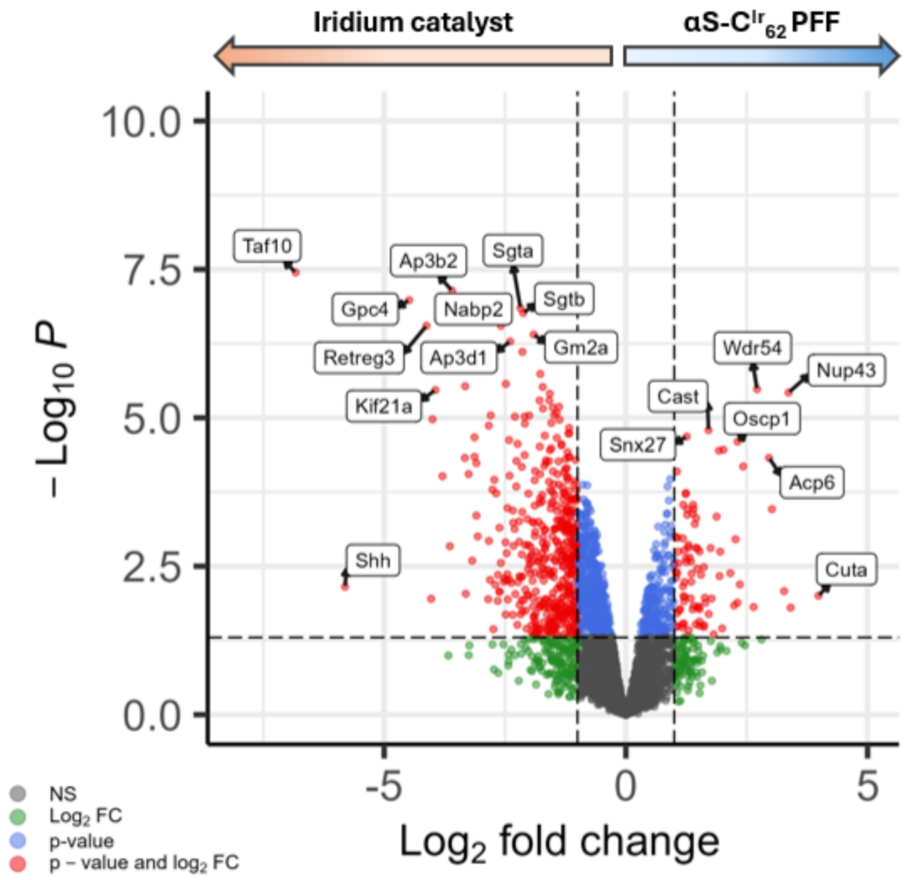


**Figure S19.** Volcano plot of lysate proteins enriched by free Iridium catalyst versus αS-$C^{Ir}_{62}$ PFF. Horizontal axis -$Log_{10}P$ = 1.313 indicates threshold of adjusted p-value = 0.05. Labeled proteins on the top right quadrant indicate enriched lysate proteins that are both statistically significant and with a >2 fold change intensity from the promiscuous labeling of the Iridium catalyst condition.

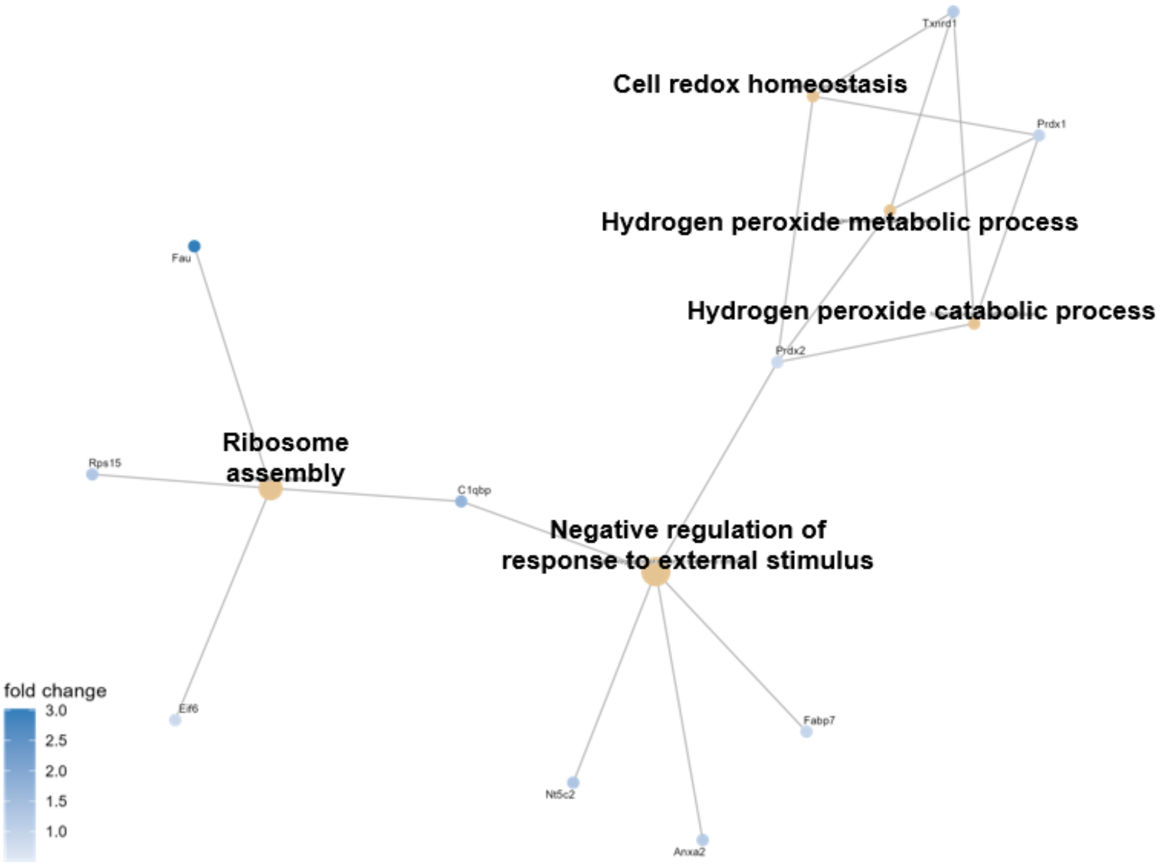


**Figure S20.** GO plot of statistically significant protein hits only from αS-$C^{Ir}_{62}$ PFF enrichment. Out of all statistically significant protein hits (adjusted p-value < 0.05) from pair-wise comparison of αS-$C^{Ir}_{62}$ PFF enriched lysate proteins versus Iridium catalyst enriched proteins, only the positive fold change terms (proteins enriched by αS-$C^{Ir}_{62}$ PFF) were taken for ORA clustering. Only blue nodes representing these αS-$C^{Ir}_{62}$ PFF enriched terms are visualized in the resultant network-node graph on a color scale based on $\log_2$ fold change.

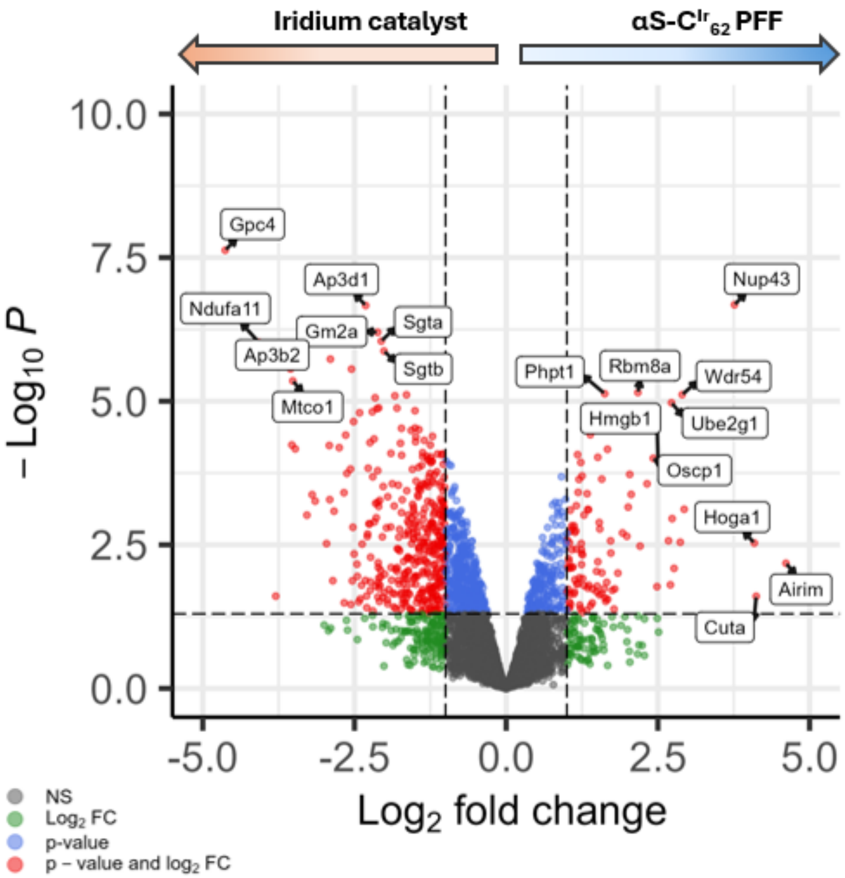


**Figure S21.** Volcano plot of lysate proteins enriched by free Iridium catalyst versus αS-$C^{Ir}_{114}$ PFF. Horizontal axis -$Log_{10}P$ = 1.313 indicates threshold of adjusted p-value = 0.05. Labeled proteins on the top right quadrant indicate enriched lysate proteins that are both statistically significant and with a >2 fold change intensity from the promiscuous labeling of the Iridium catalyst condition.

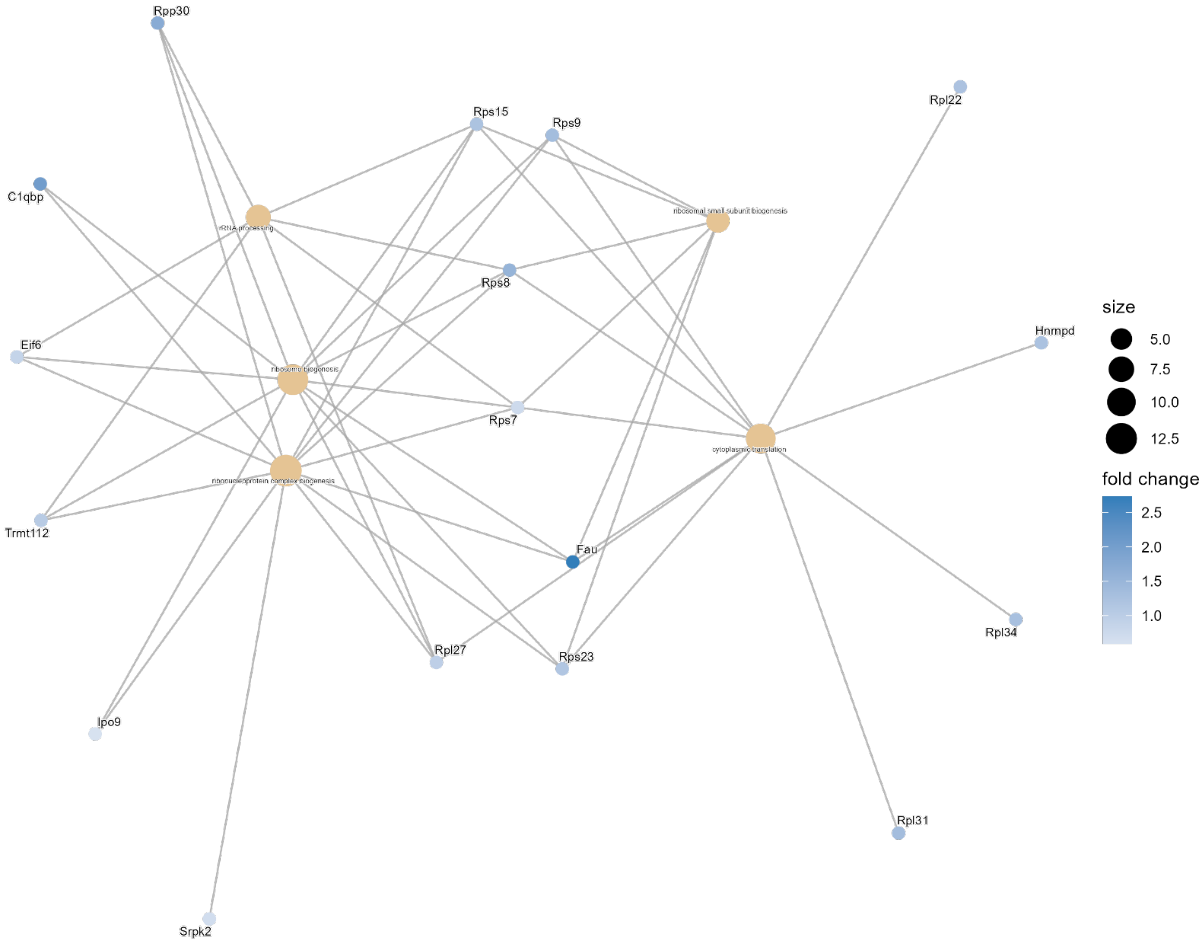


**Figure S22.** GO plot of statistically significant protein hits only from αS-$C^{Ir}_{114}$ PFF enrichment. Out of all statistically significant protein hits (adjusted p-value < 0.05) from pair-wise comparison of αS-$C^{Ir}_{114}$ PFF enriched lysate proteins versus Iridium catalyst enriched proteins, only the positive fold change terms (proteins enriched by αS-$C^{Ir}_{114}$ PFF) were taken for ORA clustering. Only blue nodes representing these αS-$C^{Ir}_{114}$ PFF enriched terms are visualized in the resultant network-node graph on a color scale based on $\log_2$ fold change

***Top 10 Enriched Proteins by Labeling Site and Conformation****

| 12 monomer | 62 monomer | 114 monomer | 12 PFF | 62 PFF | 114 PFF |
|---|---|---|---|---|---|
| Rai14 | Phactr4 | Sncb | Nup43 | Nup43 | Nup43 |
| Amot | Sncb | Nmt1 | Cast | Rbm8a | Rbm8a |
| Sprtn | Abca12 | Glud1 | Snx27 | Cbx3 | Phpt1 |
| S100b | S100b | Sprtn | Oscp1 | Phpt1 | Wdr54 |
| Actl6a | Dennd6a | Ids | Rbm8a | Oscp1 | Ube2g1 |
| Zmat2 | Tmcc2 | Ptgr3 | Ube2g1 | Snx27 | Oscp1 |
| Stk26 | Hspa4l | Supv3l1 | Acp6 | Sncb | Hmgb1 |
| Phactr4 | Clec14a | Hexb | Iah1 | Ube2g1 | Cbx3 |
| Cbln1 | Jcad | Prpf19 | Psma4 | Gbe1 | Sncb |
| Hspa4l | Amer2 | Rimbp2 | Cpne6 | Cast | Snx27 |

**ranked by significance to free Ir catalyst*

**Figure S23.** Table of the top 10 ranked enriched proteins by labeling site and conformation. Each condition was compared pairwise with the enriched interactome of free Iridium catalyst in mouse brain lysate and ranked based on adjusted p-value for fold-change intensity. The highest significantly different protein hits with respect to the Iridium control is listed above.

**Table S1.** Table of primary antibodies for ICC, Western blotting, and DNA-PAINT

| Catalogue # | Epitope | Host Species | Source | Dilution |
|---|---|---|---|---|
| 14299-1-AP | Glutamate dehydrogenase 1 (GLUD1) | Rabbit | ProteinTech | 1:10,000 (WB)<br>1:259 (DNA-PAINT) |
| | αSyn (Syn303) | Mouse | CNDR | 1:100<br>(DNA-PAINT) |
| HL2228 | S100 beta (S100β) | Rabbit | GeneTex | 1:200 (WB) |
| | p-αSyn (phosphorylated at Ser 129) (81A) | Mouse | CNDR | 1:5000 (ICC) |
| AB5543 | Microtubule associated protein (MAP2) | Chicken | EMD Millipore | 1:5000 (ICC) |
| ab24170 | Lysosomal-associated membrane protein (LAMP1) | Rabbit | Abcam | 1:1000 (ICC) |
| S32355 | Streptavidin, AlexaFluor™555 Conjugate | | Invitrogen | 1:1000 (ICC) |
| Massive-AB 2-Plex | Anti-Mouse IgG+ Docking Site 1 | | Massive Photonics | 1:100 (DNA-PAINT) |
| Massive-AB 2-Plex | Anti-Rabbit IgG+ Docking Site 2 | | Massive Photonics | 1:100 (DNA-PAINT) |
| Massive-AB 2-Plex | Imager oligo for Docking site 1+ Cy3B or ATTO655 | N/A | Massive Photonics | 0.5 nM (DNA-PAINT) |
| Massive-AB 2-Plex | Imager oligo for Docking site 2+ Cy3B or ATTO655 | N/A | Massive Photonics | 0.5 nM (DNA-PAINT) |

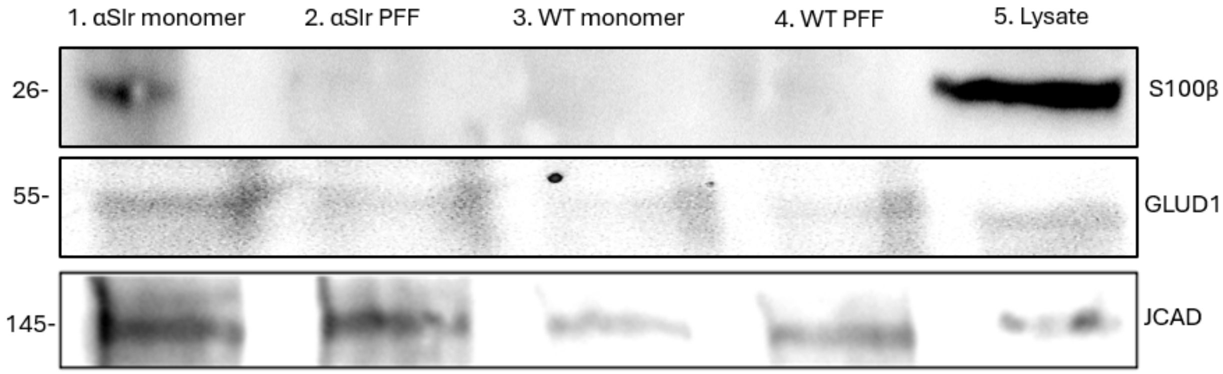


**Figure S24.** Mouse brain lysate spiked with αSIr mono or PFF (Lane 1, 2) or WT mono or PFF (Lane 3, 4) and irradiated at 445nm with Dz-BTN probe. Enriched lysate proteins were captured with Streptavidin Mag-Bead. Subsequent blot of bead eluent shown for proteomic hits S100β, GLUD1, and JCAD with non-irradiated mouse brain lysate with no αSIr treatment as positive control (Lane 5).

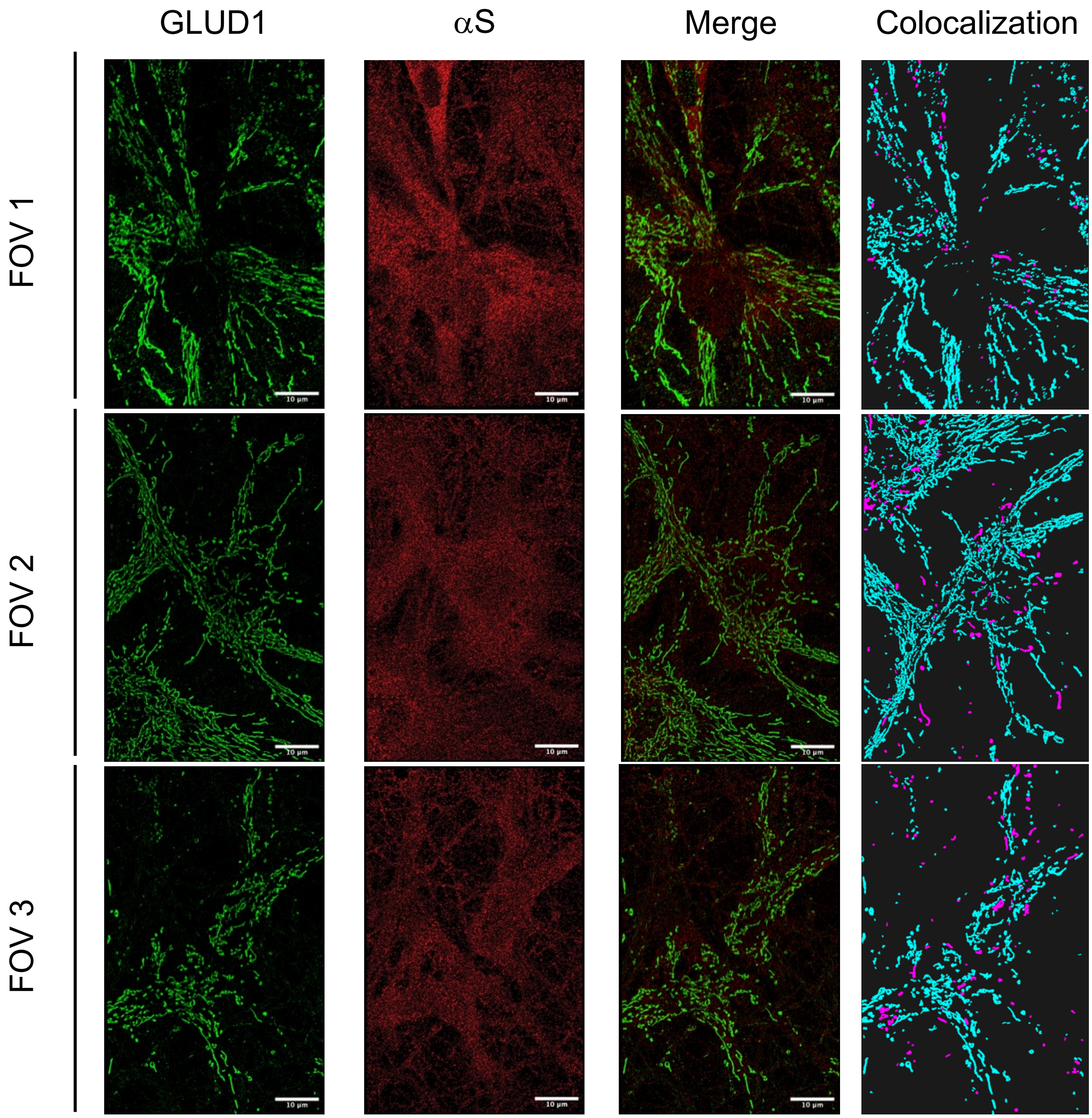


**Figure S25.** Super resolution microscopy DNA-PAINT of primary hippocampal neurons with endogenous αS signal (red) overlayed with GLUD1 signal (green). In the Colocalization images, magenta GLUD1 clusters are enriched with αS signal by a threshold of αS localization density on top and around the clusters greater than 3 standard deviations of the background αS localization. Cyan GLUD1 clusters are below this threshold and considered not to be colocalized. Primary hippocampal neurons were grown until 7 days *in vitro* after which they were fixed, probed with antibodies, and incubated with DNA-PAINT oligos before imaging. Enrichment analysis shows an average of 18.2% of all mitochondrial structures labeled by GLUD1 signal enriched by αS signal. All scale bars are 10 μM.

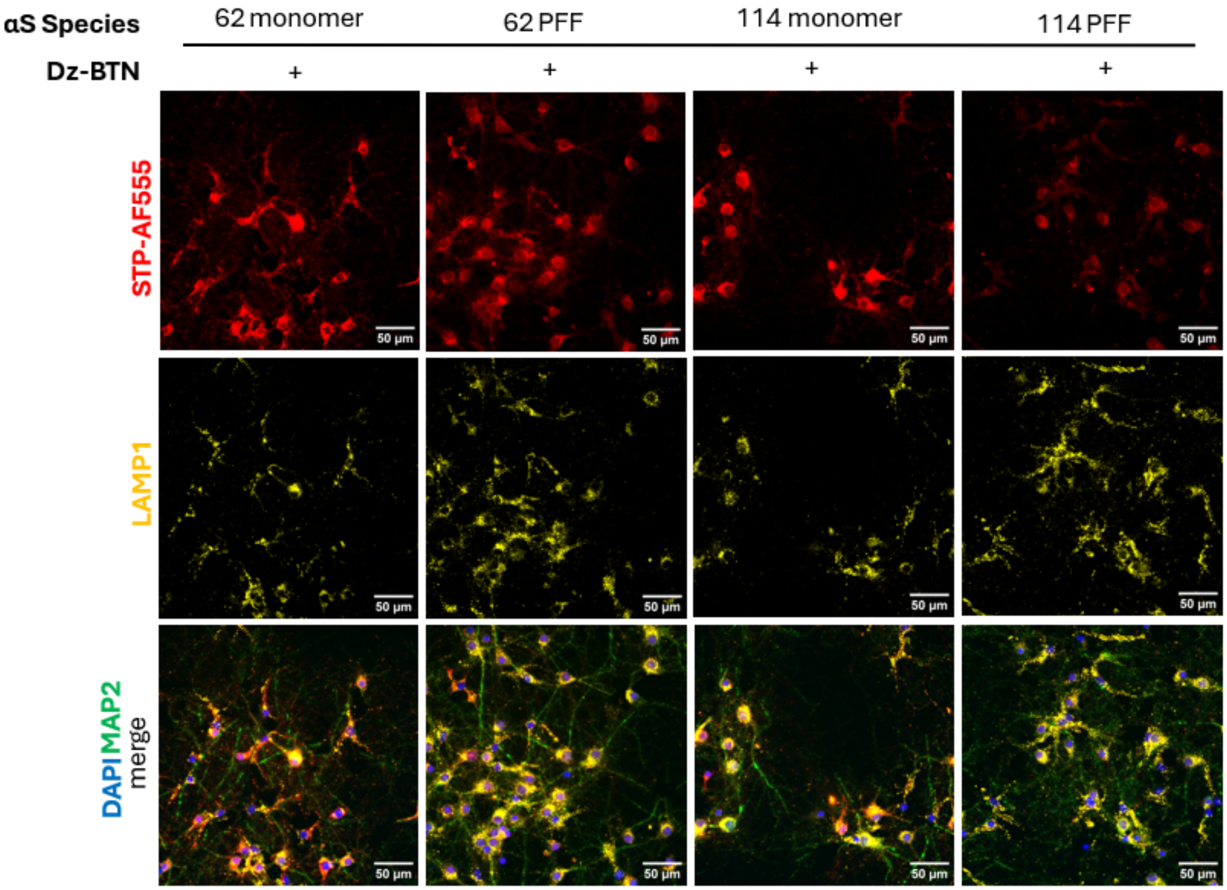


**Figure S26.** Primary hippocampal neurons treated with αS-$C^{Ir}_{62}$ and αS-$C^{Ir}_{114}$ PFF and monomer after 7 days *in vitro*. Neurons then irradiated with 445nm light with Dz-BTN, fixed, and then immunostained. Biotin-conjugated proteins were visualized with fluorescent Streptavidin-AF555 (red). Early endosomal marker LAMP1 (yellow), nuclear DAPI stain (blue), and MAP2 for neuronal processes (green) also visualized. All scale bars are 50μM.

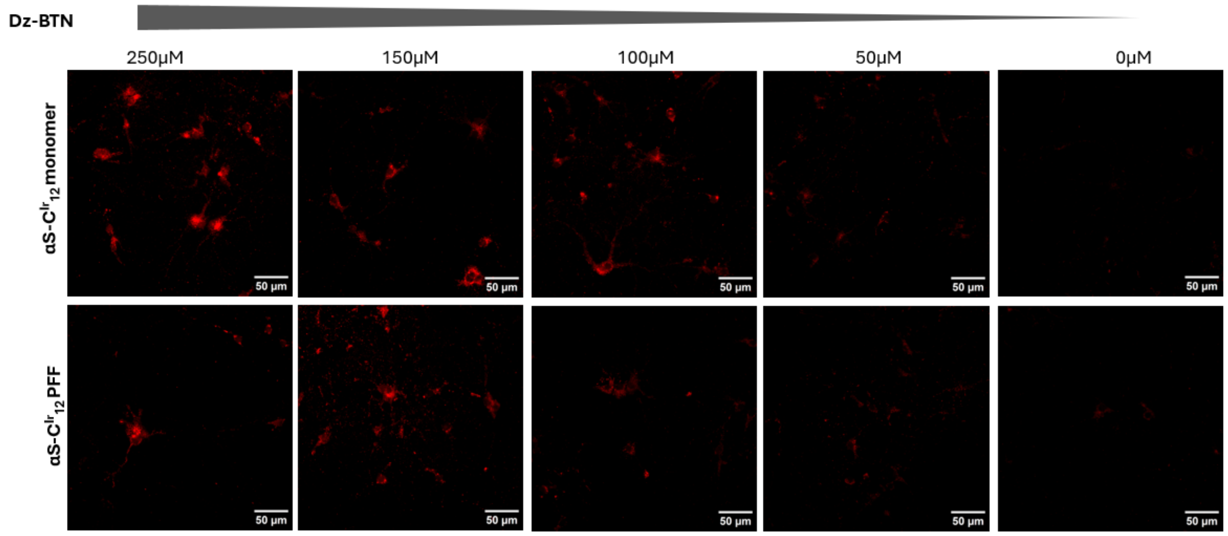


**Figure S27.** Primary hippocampal neurons treated with 47 ng/μL and 13 ng/μL of αS-$C^{Ir}_{12}$ - monomer and PFF respectively. After 24 hours of incubation, cells were treated with varying concentrations Dz-BTN and irradiated at 445nm. Photoproximity-labeled biotinylated proteins were then visualized with Streptavidin-AF555 (red). All scale bars are 50μM.

•

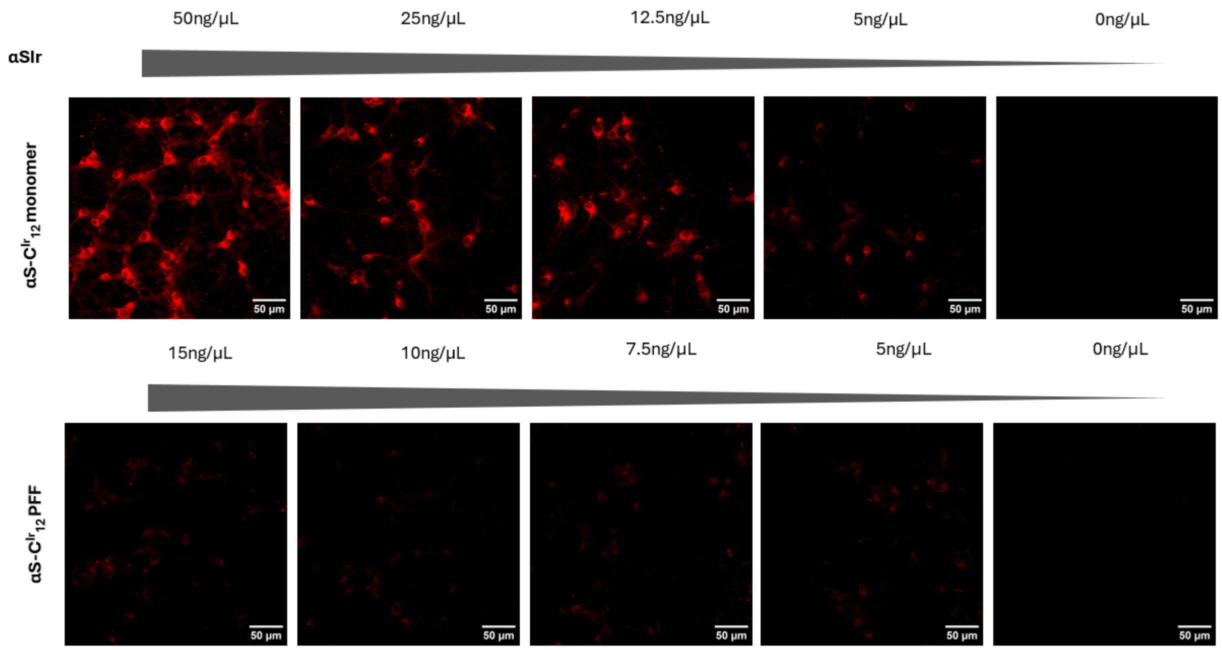


**Figure S28.** Primary hippocampal neurons treated with varying concentrations of either αS-$C^{Ir}_{12}$ monomer or PFF after which the neurons were irradiated at 445nm along with Dz-BTN. Biotin-conjugated proteins were visualized with fluorescent Streptavidin-AF555 (red). All scale bars are 50µM.

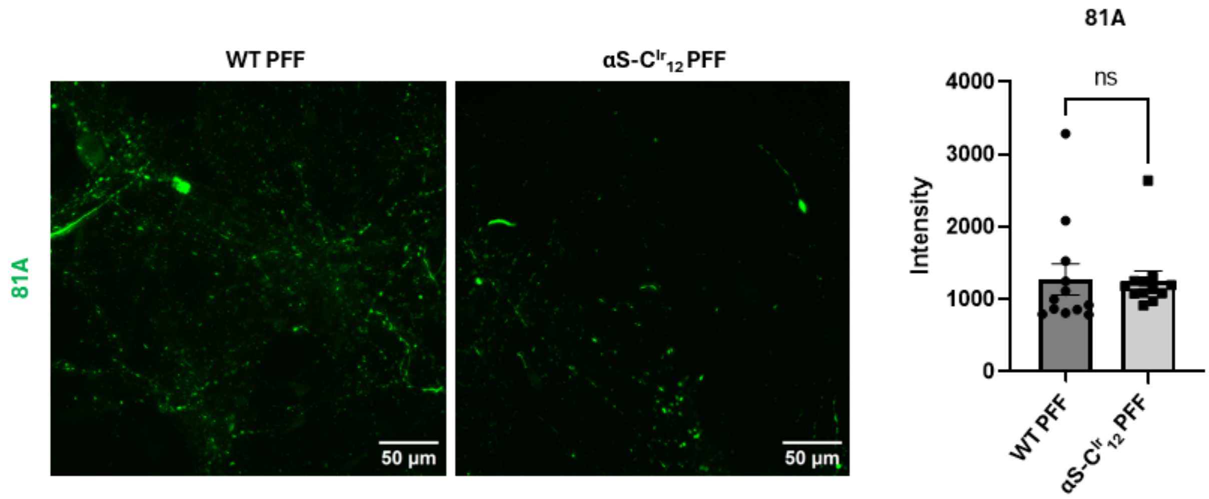


**Figure S29.** Primary hippocampal neurons treated with either 50 ng of WT αS PFF or αS-$C^{Ir}_{12}$ PFF. After 14 days *in vitro*, PFF-induced aggregation was measured using 81A intensity for pS129 (green). A two-tailed Student's t-test was used to determine significance between the WT αS seeded and αS-$C^{Ir}_{12}$ PFF seeded conditions.
